# ZEB1-dependent modulation of fibroblast polarization governs inflammation and immune checkpoint blockade sensitivity in colorectal cancer

**DOI:** 10.1101/2023.03.28.534565

**Authors:** Harald Schuhwerk, Constantin Menche, Isabell Armstark, Pooja Gupta, Kathrin Fuchs, Ruthger van Roey, Mohammed H. Mosa, Carol I. Geppert, Stefanie Bärthel, Dieter Saur, Florian R. Greten, Simone Brabletz, Thomas Brabletz, Henner F. Farin, Marc P. Stemmler

## Abstract

The EMT-transcription factor ZEB1 is heterogeneously expressed in tumor cells and in cancer-associated fibroblasts (CAFs) in colorectal cancer (CRC). While ZEB1 in tumor cells regulates metastasis and therapy resistance, its role in CAFs is largely unknown. Combining fibroblast-specific *Zeb1* deletion with immunocompetent mouse models of CRC, we observe that inflammation-driven tumorigenesis is accelerated, whereas invasion and metastasis in sporadic cancers is reduced upon fibroblast-specific loss of *Zeb1*. Single-cell transcriptomics, histological and *in vitro* characterization reveal a crucial role in CAF polarization, promoting myofibroblastic features whilst restricting inflammatory activation. *Zeb1* deficiency impairs collagen deposition and CAF barrier function but increases cytokine production, jointly promoting lymphocyte recruitment and immune checkpoint activation. Strikingly, the *Zeb1*-deficient CAF repertoire sensitizes to immune checkpoint inhibition, pointing to a therapeutic opportunity of targeting ZEB1 in CAFs and its usage as a prognostic biomarker. Collectively, we demonstrate that ZEB1-dependent plasticity of CAFs suppresses anti-tumor immunity and promotes metastasis.

## Introduction

Colorectal cancer (CRC) is the third most frequent tumor type that accounts for the second highest cancer related mortality worldwide ^1^. Despite improved diagnosis and treatment options in early stages, advanced stage CRC frequently leads to fatal metastatic relapse. Molecular classification has shown that the stroma-rich consensus molecular subtype 4 (CMS4) is linked with worst prognosis in patients ^2^. A recent refinement of this classification based on scRNA-Seq data demonstrated that fibroblast enrichment together with tumor cell intrinsic features contribute to the poor prognosis ^3^. These analyses identify cancer-associated fibroblasts (CAFs) as crucial players in the tumor microenvironment (TME) that drive disease progression, therapy resistance and metastasis ^4-7^.

Immune checkpoint blockade (ICB) therapy has shown impressive efficacy for patients with microsatellite instable (MSI) tumors ^8, 9^. These tumors display an elevated tumor mutation burden resulting in high immunogenicity. However, for most patients that present microsatellite stable (MSS) tumors, no immunotherapies are available. In preclinical mouse models the inhibition of TGFβ signaling in CAFs has been shown to promote lymphocyte infiltration and increase the efficiency of checkpoint inhibitors ^10^. Such examples highlight the concept of a stroma-directed therapy to render MSS tumors more susceptible to ICB. However, identifying specific targets to alter the fibroblast-rich stroma are needed.

Recent single cell analyses have shown that CAFs represent a heterogeneous cell population with several prototypic subtypes that were initially identified in PDAC and later in many other entities including CRC ^11-17^. Myofibroblast-like myCAFs which are in close proximity to tumor cells, secrete ECM encapsulating tumor cells, whereas iCAFs with an inflammatory expression profile are localized more in the tumor periphery and create a proinflammatory milieu ^6, 16^. As one example for the importance of CAF plasticity it has been recently demonstrated in rectal cancer that IL1 induced iCAFs adapt a senescent phenotype upon radiotherapy, which mediates radioresistance and disease progression. Therapeutic inhibition of IL1 signaling allowed to restore radiosensitivity highlighting iCAFs as promising putative therapeutic target^18^.

ZEB1 is a core EMT transcription factor that is frequently upregulated at the invasive front of CRC and other cancers, where it orchestrates tumor stemness, dissemination, metastasis and therapy resistance ^19, 20^. Its tumor cell-specific loss leads to profound changes in gene expression, impairing cell plasticity ^21-24^. In pancreas and breast cancer, ZEB1 was found to be also upregulated in the very dysplastic fibroblast-rich stroma which was correlated with poor survival ^25, 26^. However, whether ZEB1 expression contributes to CAF differentiation and function to regulate tumor progression is yet unknown. Here, we studied the role of ZEB1 in fibroblasts using mouse models for colitis-associated cancer and sporadic metastatic CRC demonstrating a key role for CAF diversification. Analysis of the tumor-immune microenvironment identified a stage-specific influence of CAF subtypes on CRC progression. Genetic inactivation of *Zeb1* reduced liver metastasis and augmented responsiveness to immune checkpoint blockade, highlighting the potential of a CAF-directed therapy.

## Results

### ZEB1 in fibroblasts affects CRC tumorigenesis in a context-dependent manner

ZEB1 is expressed heterogeneously in the TME of both human and murine tumors (Fig. 1A, B) and high expression of ZEB1 in stromal cells of pancreatic cancers is a prognostic factor for poor survival in patients ^25^. To investigate a functional involvement, we generated fibroblast-specific *Zeb1* deleted mice (Fib^ΔZeb1^), by combining the *Zeb1^fl/fl^* genotype ^27^ with the constitutive Col6a1-Cre transgene ^28^ or with the tamoxifen-inducible Col1a2-CreERT2 knock-in allele by mating ^29^. Cre recombinase activity in both lines was restricted to fibroblasts as identified by Cre reporter alleles (Extended Data Fig. 1A, B) and deletion of *Zeb1* was confirmed at the genomic (Extended Data Fig. 1C) and protein level in the colon of both models (Extended Data Fig. 1D, E). Of note, loss of *Zeb1* in fibroblasts did not compromise intestinal morphogenesis or homeostasis (Extended Data Fig. 1F, G).

**Fig. 1.**
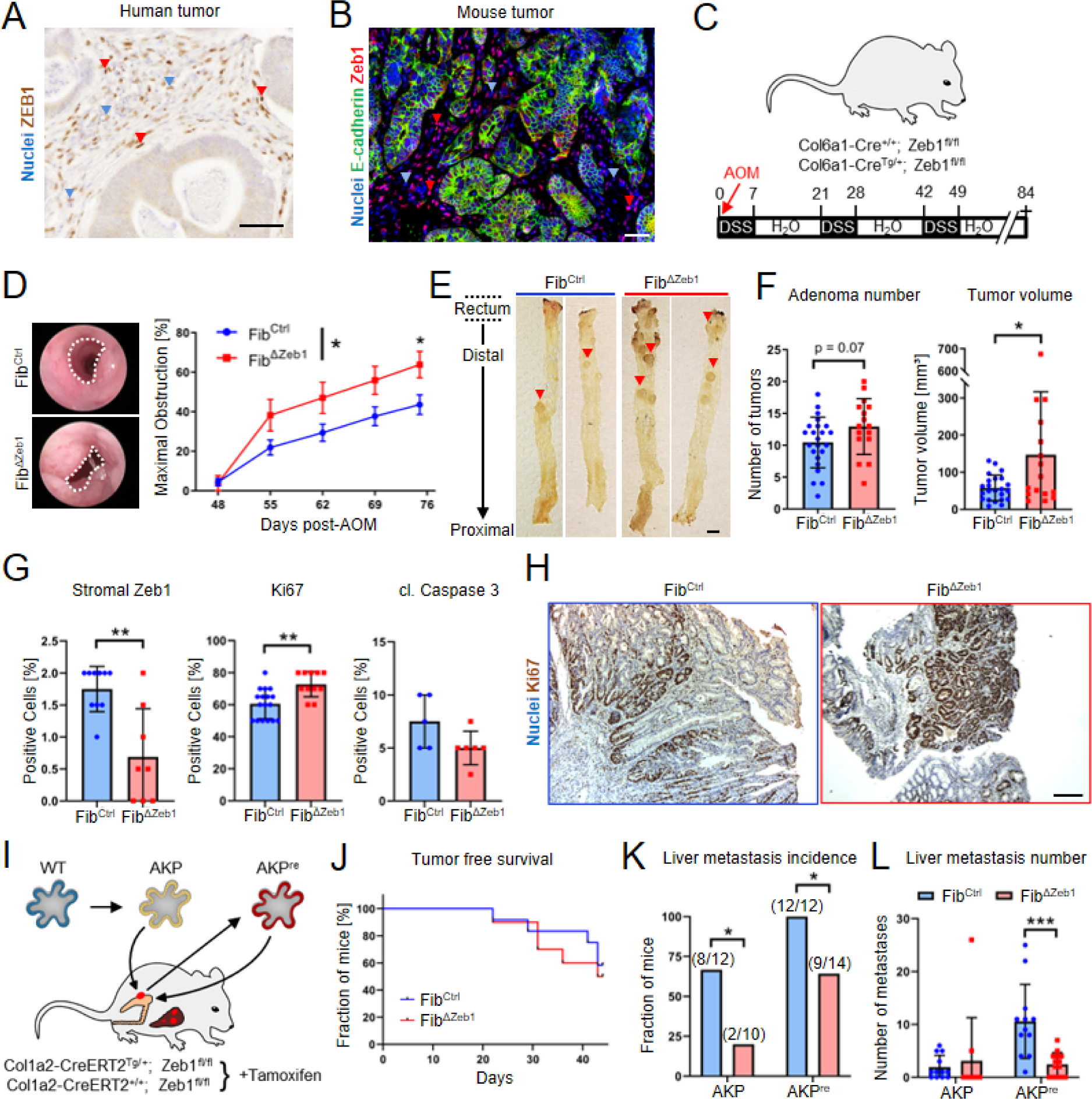
Context-dependent role of stromal ZEB1 during colorectal carcinogenesis. (A, B) anti-ZEB1 immunohistochemistry on human (A) and anti-ZEB1/E-cadherin immunofluorescence stainings on mouse (B) CRC samples. Red and blue arrowheads depict cells with high and low/absent ZEB1 detection, respectively. Scale bars: 50 µm. (C-H) AOM/DSS model (C) showing representative endoscopic images (D) (dotted lines indicate unobstructed area) and quantification of colon obstruction in Fib^Ctrl^ and Fib^ΔZeb1^ mice (n=24/17 for Fib^Ctrl^/Fib^ΔZeb1^, mean ± SD, *: p < 0.05, 2way ANOVA), longitudinal opened colons with macroscopic tumors (E) (arrowheads point out individual prominent tumors) and quantification of tumor numbers and volume (F) (n=23/16 for Fib^Ctrl^/Fib^ΔZeb1^, mean ± SD, *: p < 0.05, student’s t-test). Scale bar: 5 mm. Quantitative analysis of ZEB1 depletion, proliferation (Ki67) and apoptosis (cl. Caspase 3) (G) with representative Ki67 IHC is given (H). For ZEB1, the fraction of positive stromal cells was quantified. For Ki67 and cl. Caspase 3, the fraction of all tumor cells was quantified (n=17/11 for Fib^Ctrl^/Fib^ΔZeb1^, mean ± SD, **: p < 0.01, student’s t-test). Scale bar: 100 µm. (I-L) Orthotopic transplantation model providing an overview of generation and transplantation of tumor organoids into the caecum of Fib^Ctrl^ and Fib^ΔZeb1^ mice (I). AKP^re^ organoids were generated after re-culturing of orthotopic AKP tumor before experimental transplantation. Analysis of tumor onset (detected by palpation) after orthotopic AKP organoid transplantation (J) (n=12/10 for Fib^Ctrl^/Fib^ΔZeb1^, ns by Mantel-Cox test) as well as of liver metastasis incidence (K) (* < 0.05, Fisher’s exact test) and numbers (L) after orthotopic transplantation of AKP or AKP^re^ tumor organoids (n=12/10 for Fib^Ctrl^/Fib^ΔZeb1^ with AKP and 12/14 for Fib^Ctrl^/Fib^ΔZeb1^ with AKP^re^, mean ± SD, ***: p < 0.001, 2way ANOVA) in Fib^Ctrl^ and Fib^ΔZeb1^ mice. All mice with AKP^re^ transplantation were treated with control IgG.

Fib^ΔZeb1^ mice were subjected to the inflammation-driven AOM/DSS model ^30^ (Fig. 1C), where deletion of *Zeb1* in fibroblasts did not affect overall or tumor-free survival of mice (Extended Data Fig. 2A, B). Intriguingly, endoscopic scoring revealed increased colonic obstruction (Fig. 1D), which was confirmed by detection of bigger adenomas and a trend to increased tumor numbers in Fib^ΔZeb1^ mice at endpoints (Fig. 1E, F). Tumor differentiation was not affected by loss of stromal *Zeb1* (Extended Data Fig. 2C). However, fewer ZEB1-positive stromal cells and increased epithelial proliferation accompanied by reduced cell death was observed in Fib^ΔZeb1^ mice (Fig. 1G, H), while colonic inflammation was slightly increased (Extended Data Fig. 2D-F). These data demonstrate that loss of *Zeb1* in fibroblast promotes inflammation-driven adenoma growth.

**Fig. 2.**
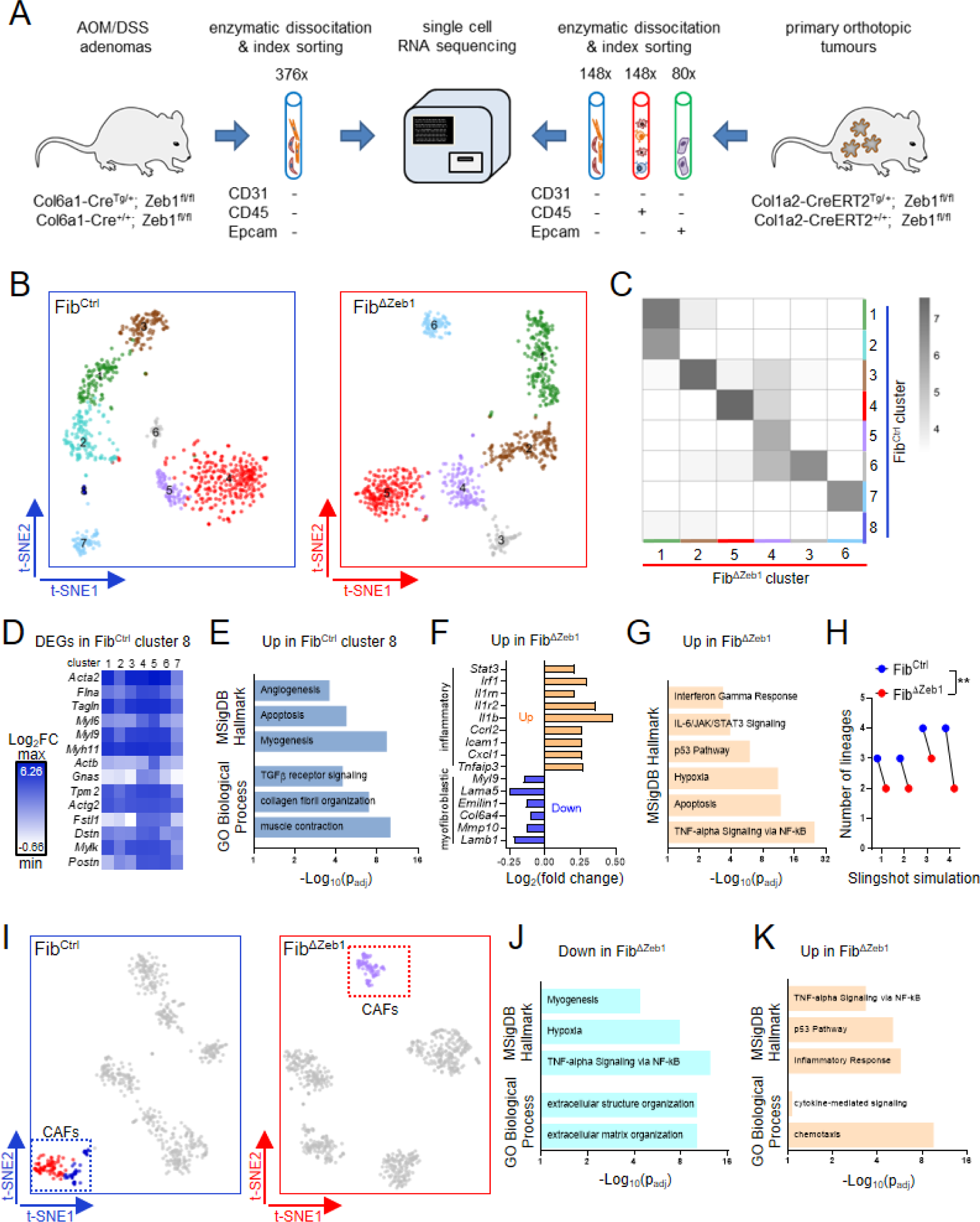
scRNA sequencing of CAFs reveals a key role of ZEB1 for fibroblast plasticity. (A) Scheme of the experimental setup for isolation of CAFs and other cell types. AOM/DSS adenomas (left) and primary tumors from the orthotopic AKP model (right) of Fib^Ctrl^ and Fib^ΔZeb1^ mice were enzymatically dissociated and fibroblasts were enriched by depletion of CD31^pos^, CD45^pos^ and Epcam^pos^ cells by flow cytometry. Epithelial and immune cells were enriched by Epcam or CD45 staining, respectively. Analyzed cell numbers per tumor are shown. Fibroblast analysis from the AOM/DSS model is shown in (B-H), while scRNA sequencing of fibroblasts, immune cells and tumor cells after orthotopic transplantation of AKP organoids is shown in (I-K). (B) Cells from n=3 mice per genotype were subjected to scRNA sequencing and t-SNE clustering independently in Fib^Ctrl^ (left) and Fib^ΔZeb1^ mice (right). (C) Reference-based annotation of fibroblast clusters of Fib^ΔZeb1^ mice using the single-cell gene expression profiles of Fib^Ctrl^ clusters as reference with the heatmap showing the distribution of cells across Fib^ΔZeb1^and Fib^Ctrl^ clusters based on ‘SingleR’ scores ^72^. Color scale in the heatmap shows the log-transformed number of cells across clusters. (D) Heatmap showing the top upregulated genes in Fib^Ctrl^ cluster 8 (FDR<0.05) as compared with the remaining Fib^Ctrl^ clusters. (E) Gene set enrichment analysis of genes significantly enriched in Fib^Ctrl^ cluster 8 compared to remaining Fib^Ctrl^ clusters. (F) Selected ‘ECM & myofibroblastic’ and ‘inflammatory’ genes show differential expression in Fib^ΔZeb1^ versus Fib^Ctrl^ cells (p-value<0.01; FDR<0.1). (G) Combined gene set enrichment analysis of all CAFs in both genotypes. (H) Slingshot analysis shows reduced number of fibroblast lineages in Fib^ΔZeb1^, independent of simulation parameters. (I) Cells from n=4 mice per genotype of the orthotopic model were subjected to scRNA sequencing and t-SNE clustering in Fib^Ctrl^ (left) or Fib^ΔZeb1^ tumors (right). Down-(J) and upregulated (K) gene sets in Fib^ΔZeb1^ CAFs compared to all Fib^Ctrl^ CAFs.

Tumor progression of sporadic CRC was modelled by orthotopic transplantation of tumor organoids ^31^ into Fib^ΔZeb1^ and Fib^Ctrl^ mice (Fig. 1I). Tumor organoids were genetically engineered from normal colonic organoids to harbor mutations in *Apc*, *Tp53* and *Kras* loci (*Apc^Δ/Δ^*, *Kras^G12D^*, *Tp53^Δ/Δ^*; AKP) and either transplanted directly into syngeneic mice, or re-cultured upon one round of orthotopic growth to promote further tumor progression *in vivo* (AKP^re^). AKP^re^ tumors indeed displayed more aggressive tumor growth with earlier onset and less differentiated morphology. However, deletion of *Zeb1* in fibroblasts did not affect engraftment of organoids, overall survival of mice, primary tumor size and morphology upon transplantation of AKP and AKP^re^ organoids (Fig. 1J and Extended Data Fig. 3A-H). Strikingly, spontaneous metastasis to the liver was decreased in Fib^ΔZeb1^ mice, regardless of AKP and AKP^re^ transplantation, as reflected in the fraction of metastasis-bearing mice and in the number of metastases per mouse (Fig. 1K, L), indicating a pro-metastatic role of ZEB1 in CRC CAFs. Overall, these findings suggest that ZEB1 in fibroblasts regulates colon cancer initiation and progression in a tumor context or stage-dependent manner.

**Fig. 3.**
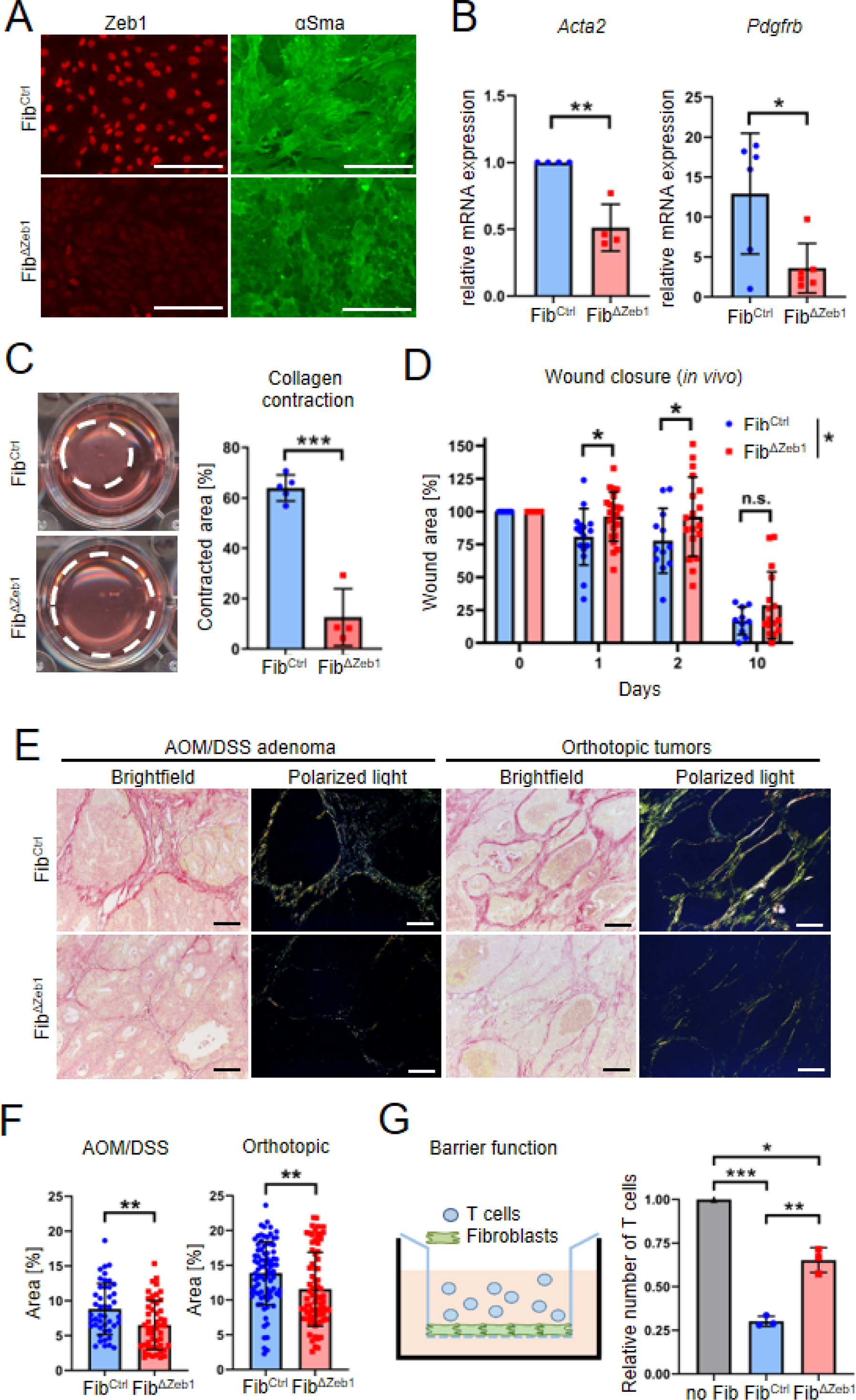
ZEB1 is critically involved in myofibroblast differentiation and functionality. (A) Representative immunofluorescence staining of ZEB1 and αSma in fibroblasts after *in vitro* recombination. Scale bars: 200 µm. (B) qRT-PCR analysis of myofibroblast marker genes *Acta2* (n=4) and *Pdgfrb* (n=6) in fibroblasts after *in vitro* recombination of *Zeb1*, mean ± SD, *: p < 0.05, **: p < 0.01, student’s t-test). (C) Representative images and quantification of collagen contraction assay after *in vitro* recombination of *Zeb1* (mean ± SD, ***: p < 0.001, student’s t-test). (D) Quantification of relative wound area in Fib^Ctrl^ and Fib^ΔZeb1^ mice during a skin wound healing model (n=16/22), mean ± SD, *: < 0.05, Holm-Šídák’s multiple comparisons test). (E-F) Representative images (E) and quantification (F) of picrosirius red staining of tumor sections from Fib^Ctrl^ and Fib^ΔZeb1^ mice after AOM/DSS and orthotopic transplantation (45/50 image sets derived from n=13/9 Fib^Ctrl^/Fib^ΔZeb1^ mice for AOM/DSS and 84/76 image sets derived from n=8/9 Fib^Ctrl^/Fib^ΔZeb1^ mice for the orthotopic tumor model, mean ± SD, **: p < 0.01, student’s t-test). Scale bar 80 µm. (G-H) Quantification of T cell migration through a transwell insert alone or with a layer of fibroblasts after *in vitro* recombination of *Zeb1* (n= 3 independent lines, mean ± SD, *: p < 0.05, **: p < 0.01, ***: p < 0.001, one-way ANOVA).

### Fibroblast diversity is reduced upon deletion of *Zeb1*

We applied scRNA-seq to gain insights into the cellular heterogeneity of CAFs and to determine the transcriptional changes upon *Zeb1* loss. AOM/DSS tumors were enzymatically dissociated and CAFs were enriched by fluorescence activated cell sorting (FACS) of CD326 (Epcam)^neg^, CD45^neg^, CD31^neg^ cells before sequencing (SORT-seq, Fig. 2A, left). Separated analysis resulted in 8 and 6 CAF clusters in Fib^Ctrl^ and Fib^ΔZeb1^ tumors, respectively (Fig. 2B). Cross-correlation of transcriptomes indicated ambiguity in Fib^ΔZeb1^ cluster number 4, as well as high similarities between Fib^ΔZeb1^ cluster 1 and Fib^Ctrl^ clusters 1 and 2. Furthermore, Fib^Ctrl^ cluster 8 was absent in *Zeb1*-deficient CAFs (Fig. 2C), altogether suggesting impaired diversification and subtype specification in Fib^ΔZeb1^ CAFs. Interestingly, the Fib^Ctrl^ cluster 8 showed elevated expression of collagens, actin cytoskeletal and ECM-related genes that contributed to enrichment of related expression signatures (Fig. 2D, E). These findings indicate reduced myofibroblast polarization upon *Zeb1* loss. Consistently, global comparison between Fib^ΔZeb1^ and Fib^Ctrl^ CAFs showed downregulation of genes involved in ECM organization (e.g. *Lamb1*, *Mmp10*, *Col6a4*, *Emilin1*, *Lama5* and *Myl9* (Fig. 2F). On the contrary, inflammatory signatures, including genes such as *Tnfaip3*, *Cxcl1*, *Icam1*, *Ccrl2*, *Nfkbia*, *Il1b* and *Irf1* were upregulated in Fib^ΔZeb1^ CAFs (Fig. 2F, G). These data show that *Zeb1*-deficient AOM/DSS CAFs exhibit enhanced inflammatory and reduced myofibroblastic features. Supporting this notion, slingshot analyses showed a reduced number of fibroblast trajectories in Fib^ΔZeb1^ tumors (Fig 2H).

We also examined non-inflammation-driven orthotopic tumors by scRNAseq. Following FACS, CAFs were analyzed in parallel to isolated CD45^pos^ immune and Epcam^pos^ tumor cells (Fig. 2A, right). Clustering analysis matched the isolated cell lineages and resulted in two CAF clusters for Fib^Ctrl^ but only one for Fib^ΔZeb1^ tumors (Fig. 2I). Separated clustering of CAFs confirmed the previous notion that loss of *Zeb1* limits the CAF repertoire (Extended Data Fig. 4A). The transcriptomes of Fib^Ctrl^ CAFs correlated well with “iCAF” and “myCAF” archetypes described in pancreatic ductal adenocarcinoma ^13, 16^, whereas Fib^ΔZeb1^ CAFs displayed more ambiguity. Cross-annotation with Fib^Ctrl^ CAFs identified the gene set of the myofibroblastic cluster 5 to be most underrepresented upon loss of *Zeb1* (Extended Data Fig. 4B, C). Consistent with AOM/DSS CAFs, differential gene expression analysis between Fib^Ctrl^ and Fib^ΔZeb1^ CAFs showed reduced signatures for ECM organization and increased inflammatory terms (Fig. 2J, K). In summary, our data show that independent of the CRC model, CAF diversification and specification is strongly impaired when ZEB1 is absent.

**Fig 4.**
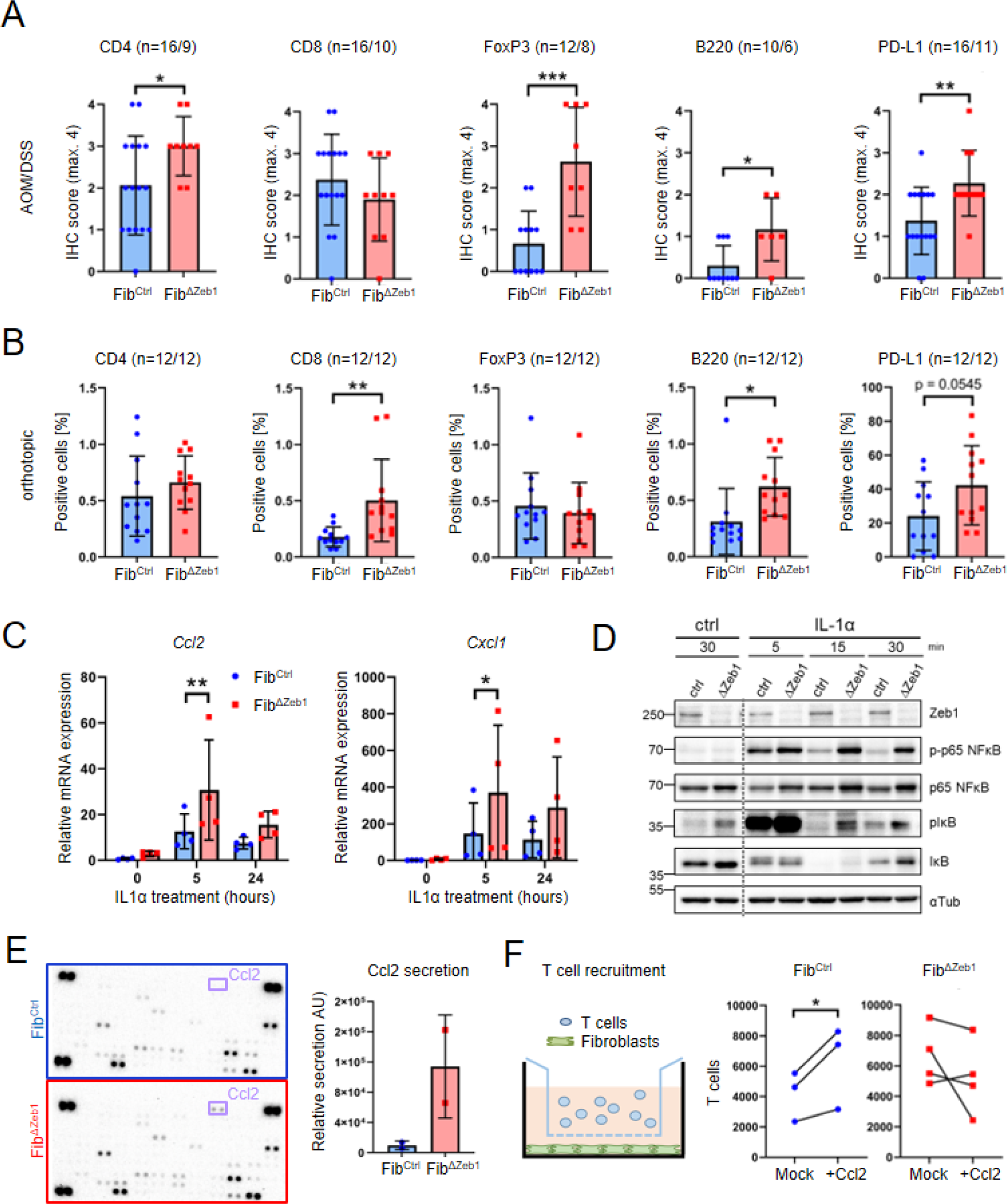
ZEB1 attenuates inflammatory signaling in fibroblasts and limits immune cell infiltration in multiple CRC models. (A-B) IHC-based quantification of immune cell infiltration and PD-L1 expression of tumors from Fib^Ctrl^ and Fib^ΔZeb1^ mice in the AOM/DSS model (A) and after orthotopic transplantation of AKP tumor organoids (B). Numbers of experimental mice per genotype are indicated (mean ± SD, *: p < 0.05, **: p < 0.01, ***: p < 0.001, student’s t-test). (C) qRT-PCR analysis of *Cxcl1* and *Ccl2* mRNA expression in fibroblasts after *in vitro* recombination of *Zeb1* and stimulation with IL1α (n=4, mean ± SD, *: p < 0.05, **: p < 0.01, 2way ANOVA). (D) Western blot of NFκB pathway activity in fibroblasts after 15 min of IL1α stimulation. α-Tubulin detection was used as a loading control. (E) Secretome analysis of Fib^Ctrl^ and Fib^ΔZeb1^ CAFs isolated from AOM/DSS tumors using a secretome array showing one representative exposure of the arrayed membrane (left) and quantification using two independent pairs of CAFs (right). (F) Quantification of T cell attraction to fibroblasts after *in vitro* recombination of *Zeb1* without (Mock) or with (+Ccl2) addition of exogenous Ccl2 (n=3/4, Fib^Ctrl^/Fib^ΔZeb1^ lines, *: p < 0.05, paired t-test).

### Primary function of ZEB1 for myofibroblast differentiation

We next wanted to gain mechanistic insight how ZEB1 regulates CAF subtypes and polarization. We established adherent colon fibroblast cultures from *Zeb1^fl/fl^* mice, which showed a typical myofibroblast-like morphology *in vitro*. Lentiviral transduction of Cre recombinase resulted in efficient *Zeb1* deletion, which was confirmed by immunofluorescence staining (Fig. 3A). Of note, *Zeb1* deficiency in these fibroblasts resulted in a loss of fibrillary αSma staining, a well-recognized marker for myofibroblast differentiation and CAF activation^6^. Consistently, *Zeb1*-deleted fibroblasts showed lower mRNA abundance of myofibroblast markers (Fig. 3B) and strongly reduced collagen contraction (Fig. 3C), altogether indicating a requirement of *Zeb1* for myCAF polarization and function. To study myofibroblast-specific functions *in vivo*, skin wound healing assays revealed delayed wound closure in Fib^ΔZeb1^ compared to Fib^Ctrl^ mice (Fig. 3D). Furthermore, the impaired myCAF polarization in AOM/DSS adenomas and primary orthotopic tumors was associated with reduced deposition of collagen (Fig. 3E, F). Since collagenous ECM in tumors can form a physical barrier for immune cell migration, we conducted an *in vitro* migration assay to test the ability of fibroblasts to block passage of mouse splenocytes (Fig. 3G). Strikingly, Fib^ΔZeb1^ fibroblasts showed a pronounced barrier defect. Collectively, these data demonstrate a critical role of ZEB1 in myofibroblast specification and function.

### Loss of *Zeb1* in fibroblasts induces a pro-inflammatory tumor microenvironment

To clarify whether the inflammatory polarization in combination with the reduced collagen deposition of *Zeb1*-deleted CAFs affects intratumor immune cell infiltration, we performed IHC analyses. Strikingly, in AOM/DSS tumors, we found increased infiltration of CD4+ T cells and FoxP3+ Tregs, but not of CD8+ cells, into Fib^ΔZeb1^ tumors (Fig. 4A), indicating modulation of adaptive anti-tumor immunity. Concomitantly, increased B cell infiltration and an overall upregulation of the immune checkpoint molecule PD-L1 was observed, suggesting tolerance induction upon T cell activation. Similar to AOM/DSS adenomas, increased T and B cell infiltration and expression of PD-L1 was observed in Fib^ΔZeb1^ orthotopic tumors (Fig. 4B). Yet in contrast to AOM/DSS tumors, particularly CD8+ T cells were enriched. Since the primary tumor size was comparable in Fib^Ctrl^ and Fib^ΔZeb1^ orthotopic tumors (Extended Data Fig. 3B, G) the observed T cell infiltration apparently did not result in an effective anti-tumor response, presumably due to partial T cell inactivity. Indeed, scRNAseq analysis of the T cell clusters showed a substantial decrease in the proliferation genes *Mki67*, *Top2a* (Topoisomerase 2) and *Ccna2* (Cyclin A2) in Fib^ΔZeb1^ compared with Fib^Ctrl^ T cells (Extended Data Fig. 5).

**Fig. 5.**
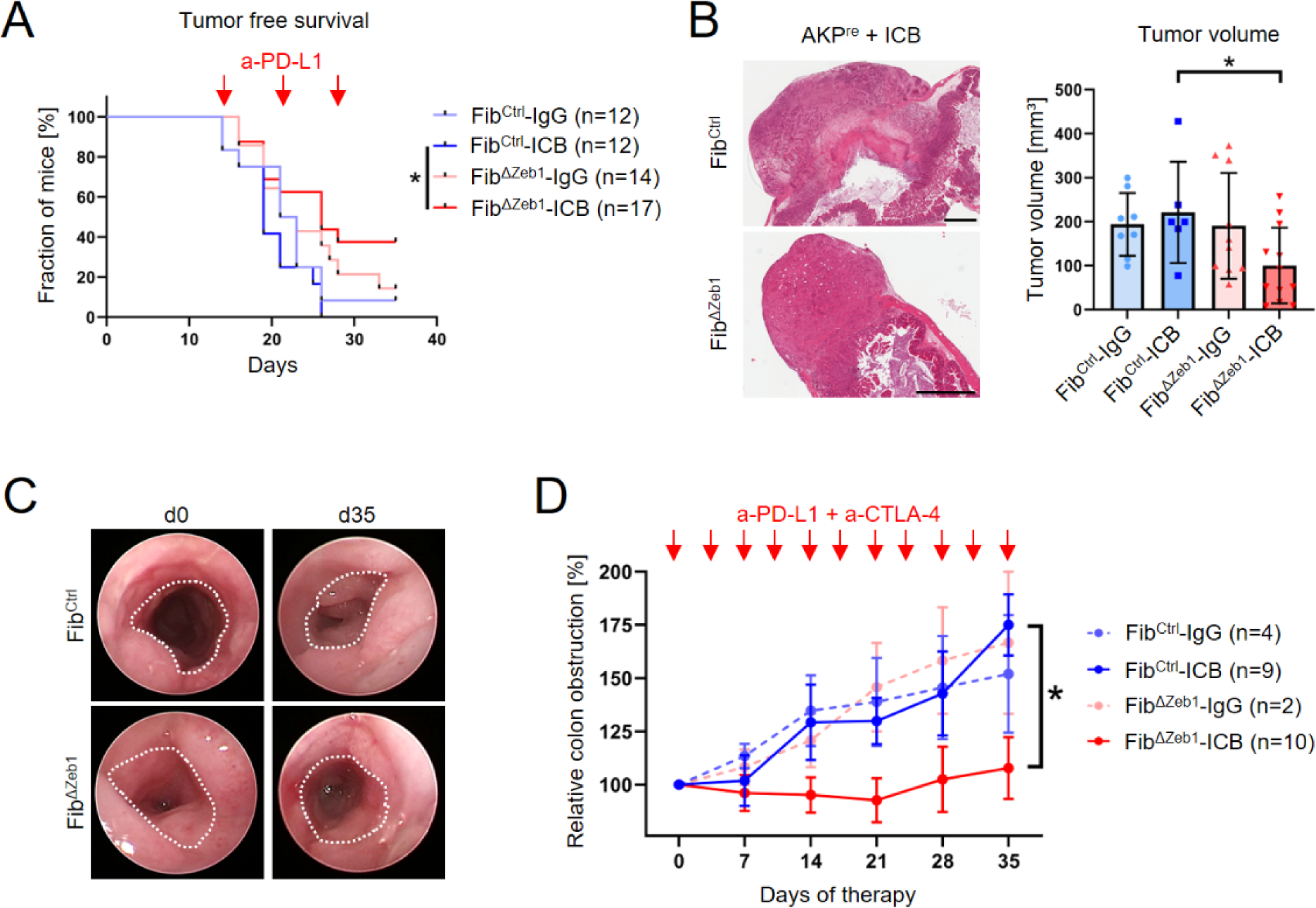
Loss of *Zeb1* in fibroblasts enables response to ICB therapy. (A) Kaplan-Meier analysis showing tumor free survival of Fib^Ctrl^ and Fib^ΔZeb1^ mice after orthotopic transplantation of AKP^re^ organoids and intraperitoneal injection of anti-(a)-PD-L1 antibodies or control IgGs, as indicated by the red arrows. Mice were considered tumor-free if no tumor was detected by palpation. Numbers of experimental mice per condition are indicated. (B) Representative H&E images and quantification of volumes of tumors after orthotopic transplantation of AKP^re^ tumor organoids and ICB. Only tumors collected after day 28 were included in this analysis (mean ± SD) and IgG-treated mice contributed to initial AKP^re^ analysis. (C, D) ICB in the AOM/DSS model, applied by two intraperitoneal injections of a-PD-L1 and a-CTLA-4 antibodies or control IgGs per week starting at day (d) 70 of AOM/DSS tumorigenesis for all mice displaying at least 25-30% of colon obstruction. This time point was set to d0 of ICB. Representative endoscopic images at d0 and d35 of ICB in the AOM/DSS model are shown. Dotted line indicates unobstructed area (C). Quantification of colon obstruction relative to d0 (D). Numbers of experimental mice per condition are indicated (mean ± SD, *: p < 0.05, 2way ANOVA with Holm-Šídák’s multiple comparisons test).

To investigate whether the differences between both models can be explained by the inflammatory environment during tumorigenesis of the AOM/DSS model, we employed another model of sporadic CRC, in which AOM-induced mutagenesis in combination with intestinal epithelial cell-specific *Tp53* deletion results in invasive autochthonous CRC (Extended Data Fig. S6A^32, 33^). Of note, deletion of *Zeb1* in fibroblasts in this model resulted in fewer, smaller and less invasive tumors, indicating delayed tumor progression in Fib^ΔZeb1^ mice (Extended Data Fig. 6B-D). Again, deletion of *Zeb1* in fibroblasts enhanced immune cell infiltration, with a comparable pattern as in the orthotopic model (Extended Data Fig. 6E). Collectively, our data show that *Zeb1* loss in CAFs in various CRC models induces a context-dependent immune cell infiltration combined with PD-L1 upregulation indicating a critical immunomodulatory role. To study the mechanism for the altered immune cell infiltration in *Zeb1*-deficient CAFs, we tested the response of cultured fibroblasts to the chemokine IL1α, a crucial stimulator of inflammatory polarization in CRC CAFs ^18^. We monitored basal and IL1α induced NFκB activation in cultured Fib^Ctrl^ and Fib^ΔZeb1^ fibroblasts by qRT-PCR and western blot analyses. Upon stimulation, Fib^ΔZeb1^ fibroblasts showed increased expression of NFκB targets including *Ccl2* and *Cxcl1* that both have been described as iCAF markers ^12, 13, 18^ (Fig. 4C). Consistently, strongly increased levels of phosphorylated NFκB and IκBα were observed in lysates of Fib^ΔZeb1^ fibroblasts that were stimulated with IL1α for 15 min (Fig. 4D), when Fib^Ctrl^ cells already partially resolved the signal. This kinetic of *Zeb1*-deficient fibroblasts suggests an overshooting activation of the pathway. Secretome analysis in resting CAFs from AOM/DSS tumors consistently identified increased basal levels of Ccl2 upon *Zeb1* loss (Fig. 4E).

Our data suggests that increased Ccl2 from *Zeb1* deficient CAFs may act as a potent lymphocyte chemoattractant. Thus, the influence of Fib^Ctrl^ or Fib^ΔZeb1^ fibroblasts on T cell recruitment was tested in a transwell migration assay with or without exogenous Ccl2 using stimulated T cells in co-culture with fibroblasts. Consistent with the increased inflammatory profile, migration of T cells towards a Fib^ΔZeb1^ fibroblast monolayer was increased in comparison to Fib^Ctrl^, albeit not significantly. However, while the attraction of T cells towards Fib^Ctrl^ CAFs was enhanced by addition of Ccl2, no additional increase was observed in Fib^ΔZeb1^ CAFs, indicating saturation by endogenous Ccl2 (Fig. 4F). Collectively, our data demonstrate that ZEB1 in fibroblasts limits tumor immune infiltration by reduced immune cell chemoattraction in addition to formation of a physical barrier.

### Loss of *Zeb1* in CAFs enhances response of CRC to immune checkpoint blockade

Given that the majority of sporadic CRCs is considered refractory to ICB therapy, the increased immune cell infiltration and induction of checkpoint molecules observed in Fib^ΔZeb1^ mice may point to a strategy to render non-hypermutated tumors sensitive. We investigated this hypothesis in orthotopic and AOM/DSS tumors. Following transplantation of AKP^re^ organoids, tumor-bearing mice were injected subcutaneously with anti-PD-L1 antibodies for 3 weeks. Strikingly, anti-PD-L1 therapy but not control IgG significantly delayed tumor growth of Fib^ΔZeb1^ in comparison with Fib^Ctrl^ mice, eventually resulting in smaller Fib^ΔZeb1^ tumors at the endpoint (Fig. 5A, B). Additionally, infiltration of immune cells was further increased by ICB and PD-L1 expression was lost in tumors from treated Fib^ΔZeb1^ mice, indicating successful reactivation of the adaptive immune response (Extended Data Fig. 7). In autochthonous AOM/DSS tumors, we employed dual ICB using anti-PD-L1 and anti-CTLA-4 antibodies, as we found a substantial increase in the number of Tregs in tumors of Fib^ΔZeb1^ mice in this model (Fig. 4A). Tumor growth was monitored longitudinally via endoscopy, and adenoma-bearing mice were subjected to ICB after recovery from cyclic DSS-induced inflammation on day 70. Strikingly, dual ICB abrogated tumor growth in Fib^ΔZeb1^ mice (Fig. 5C, D), demonstrating that loss of *Zeb1* in fibroblasts induces sensitivity to ICB in both colitis-induced and sporadic tumor models.

## Discussion

Functional diversification of CAFs is a result of a coevolution with tumor cells in the TME. Using autochthonous and organoid transplantation models of CRC, we have discovered a key role of ZEB1 in governing CAF plasticity. *Zeb1* deletion severely impairs myofibroblastic and modulates inflammatory CAF functions, jointly increasing tumor immune cell infiltration. Consequently, inflammation-driven adenoma formation is augmented, yet more progressed cancer models show reduced invasiveness and metastasis. Despite this stage-dependent outcome, common immune modulation in Fib^ΔZeb1^ tumors leads to activation of immune checkpoints. We show that targeting ZEB1 sensitizes to ICB reinforcing the translational rationale to target specific CAF subtypes for therapy of microsatellite-stable tumors.

Our data identifies that ZEB1 in CAFs acts as an immuno-suppressor that promotes malignant tumor progression. As both archetype populations, *i.e.* iCAFs and foremost myCAFs, are strongly dysregulated in the absence of ZEB1, we conclude that the general CAF plasticity and malignancy-promoting effect critically depends on ZEB1 in CRC. This is supported by the reduced number of cell clusters in scRNA data of both models, the impaired myofibroblastic functionality and augmented response to inflammatory stimuli observed *in vivo* and *in vitro*. While previous reports have established that EMT(-related-) TFs ZEB1, Snail, Twist and Prrx1 can regulate the classical mechanoinvasive features of CAFs directly affecting tumor malignancy ^34-36^, our data highlight a critical impact of CAF plasticity on modulation of anti-tumor immunity that may contribute to the poor prognosis reported in breast and pancreatic cancer patients with elevated ZEB1+ stromal cells ^25, 37^.

As a mechanistic basis for enhanced inflammation and immune cell infiltration, we show reduced collagen deposition, increased inflammatory signaling and chemoattraction in *Zeb1* deficient CAFs collectively modulating the tumor immune TME. The tumor stage-dependent consequences can be explained by the known tumor promoting role of inflammation in AOM/DSS Fib^ΔZeb1^ tumors ^7, 28, 30, 32, 38^. Consistently, we found strongly increased infiltration of B cells, CD4+ T cells and Foxp3+ Tregs in Fib^ΔZeb1^ tumors accompanied by enhanced PD-L1 and CTLA-4 abundance. Since no effect was observed on the influx of CD8+ T cells, we reason that impaired T cell effector function may have been caused by induction of an immune checkpoint involving Tregs. This is in agreement with previous reports on tolerance induction in AOM/DSS and colitis-associated CRC ^39-41^. In contrast, the increased infiltration of B cells, CD4+ and CD8+ T cells in the Fib^ΔZeb1^ non-inflammatory sporadic tumors was associated with delayed tumor progression. Taken together, we conclude that loss of *Zeb1* in CAFs facilitates inflammation, immune infiltration and co-activated immunosuppression.

We furthermore identified ZEB1 as an important regulator of balancing myofibroblastic and inflammatory functions of CRC CAFs. Of note, TGFβ signaling has been shown to be essential for acquisition of classical myofibroblast phenotypes ^6, 10, 12, 13, 42^ and in tumor cells, ZEB1 is a key mediator of TGFβ signaling ^19, 21, 27, 43^. This indicates that ZEB1 may act as a mediator of TGFβ signaling during myofibroblast polarization. Simultaneously, *Zeb1*-deleted fibroblasts consistently displayed alterations in inflammatory gene expression in tumors and increased NFκB activation in response to IL1α *in vitro*. Given that IL1α is a strong inducer of iCAFs ^6, 12, 18^, these data show that acquisition of core iCAF features is accomplished independent of ZEB1, whereas ZEB1 function is indispensable for fine-tuning of the iCAF phenotype. In this regard, it is important to mention that ZEB1 has been shown to act as a direct transcriptional (co-)inducer of inflammatory gene expression in several cell types, such as breast cancer cells and fibroblasts ^26, 44^, corneal fibroblasts, hematopoietic and myeloid cells ^45-49^. Together these data suggest that ZEB1 elicits context-dependent effects on inflammation.

The observed sensitization of tumors to ICB independent of the disease context points towards a general CAF-engaging immune checkpoint activation in CRC, in line with the refractoriness non-hypermutated tumors to ICB ^7, 42, 50^. Moreover, it highlights the concept of a stroma-directed therapeutic approach. Thus, ZEB1 expression in CAFs may serve as a negative predictive marker for ICB efficacy in CRC. Accordingly, agents that modulate CAF identities may be beneficial to enhance immune infiltration and ICB sensitivity in the clinics. For instance, inhibiting the DDR kinase ATM has recently been shown to inhibit myofibroblastic features, increase immune infiltration and additively sensitize to immune checkpoint therapy in subcutaneous tumor models ^51^. Because targeting ZEB1 as a TF is challenging, new strategies will be required. In this regard, we recently discovered an actionable vulnerability in ZEB1^high^ cancer cell sub-populations by inhibiting the DNA damage response (DDR) nuclease MRE11^43^. Along that line, such selective pharmacological approaches or development of specific PROTACs bear the promise for targeting ZEB1 in CAFs directly or indirectly. However, better understanding of CAF polarization and function is needed, especially in response to standard therapies. For instance, therapy-induced senescence in inflammatory CAFs favors therapy resistance and a poor outcome in rectal cancer ^18^. Likewise, depletion of αSMA-high myCAFs, or deletion of type 1 collagen in myofibroblasts was shown to aggravate the disease course in PDAC and experimental liver metastasis ^52-54^. Hence, targeting of regulators of CAF plasticity, such as ZEB1, rather than fully depleting integral components of the ECM or entire CAF subtypes, might be beneficial. Our study suggests that targeting ZEB1 in CAFs may turn immunologically “cold” into “hot” CRCs and thereby sensitize CRC patients to ICB. Given the known role of ZEB1 in tumor cells to induce EMT, stemness and chemoresistance, combined targeting of ZEB1 in fibroblast and tumor cells might act synergistically to improve CRC therapy.

## Methods

### Ethics statement

Animal husbandry and all experiments were performed according to the European Animal Welfare laws and guidelines. The protocols were approved by the committee on ethics of animal experiments of Bavaria (Regierung von Unterfranken, Würzburg; TS-18/14, 55.2-DMS-2532-2-270, -2-952 and -2-1133) and of Hessen, Germany (Regierungspräsidium Darmstadt F123/1031, F123/1040 and F123/2001). Power analysis was used to calculate the sample size required for animal experiments. Animals were kept on a 12:12 h light-dark cycle and provided with food and water *ad libitum* in the animal facilities of the Friedrich-Alexander University of Erlangen-Nürnberg and the Georg-Speyer-Haus Frankfurt. Col1a2-CreERT2^Tg/+^;*Zeb1^fl/fl^* and Col6a1-Cre^Tg/+^;*Zeb1^fl/fl^* mice were kept on FvB and C57BL/6 backgrounds, respectively. Col6a1-Cre^Tg/+^;*Zeb1^fl/fl^*;Vil-Flp^Tg/+^;*Tp53^FRT/FRT^* Fib^ΔZeb1^ mice were kept on a mixed FvB/C57BL/6 background.

### Mouse models of CRC

Rosa26-tdTomato ^55^ and Rosa26-mTmG ^56^ mice have been described previously. Conditional *Zeb1* knockout mice ^27^ were crossed with mice expressing Col6a1-Cre ^28^ or Col1a2-CreERT2^29^ to generate Col6a1-Cre-positive Fib^ΔZeb1^ mice or Col6a1-Cre-negative Fib^Ctrl^ mice. Age-matched littermates of both sexes were used for experiments and randomly assigned to groups within genotypes.

AOM/DSS inflammation-driven autochthonous adenoma formation was induced as described^30^. Briefly, mice were injected intraperitoneally (i.p.) with 10 mg/kg of azoxymethane (AOM, Sigma/Merck, A5486) in 0.9% NaCl), prior to administration of three cycles of 1.75% dextran sulfate sodium (DSS, MP Biomedicals, SKU-0216011080) in the drinking water *ad libitum*, each interrupted by 2 weeks of regular water and finally sacrificed at day 84 or at ethical end points. Colonic inflammation and tumor growth was monitored by endoscopy under isoflurane anesthesia employing the Colorview endoscopic system (Karl Storz), as described before ^57^, expressed as numbers of adenoma and/or the maximal colon obstruction by the adenoma in percent of the full colonic diameter at the most blocked site. Colonic inflammation was visually scored from ‘0’ (zero) to ‘4’ (four), with ‘0’ representing normal, ‘1’ signs of thickening, increased vascularity, and/or fibrin visible, moderate granularity and/or stool being still shaped, ‘2’ stronger thickening, vascularization, granularity and unshaped stool (no blood), ‘3’ loss of transparency, extreme granularity, rectal blood (clots) and/or traces of blood in the mucosa and/or in the unshaped or spread stool, and ‘4’ (partially) liquid stool, several sites of bloody mucosal damage or non-rectal liquid blood in the colon. Stool consistency was scored individually from ‘1’ to ‘4’ as well, with ‘0’ being normal and solid, ‘1’ softer than usual, ‘2’ unshaped, ‘3’ almost liquid but containing solid pieces of stool and/or containing traces of (clotted) blood, ‘4’ being liquid and/or mostly bloody. For immune checkpoint inhibition, mice were injected i.p. with anti-PD-L1 (BXC-BE0101) and anti-CTLA-4 (BXC-BE0131) antibodies or isotype controls (BXC-BE0094 and BXC-BP0087) twice per week (10 mg/kg each, Bio X cell). Therapies started at day 70 for all mice, unless colon obstruction by adenomas did not reach 25-30%, as determined by endoscopy. Adenoma growth was then monitored weekly via endoscopy.

The AOM/p53 model was based on the previously reported Tp53ΔIEC model ^32, 33^, using Villin-Flip mediated recombination and intestinal epithelium-specific inactivation of biallelic FRT-flanked *Tp53* (intron 1 and 10). Experimental animals were generated by sequential crossings to generate experimental Col6a1-Cre^Tg/+^;*Zeb1^fl/fl^*;Vil-Flp^Tg/+^;*Tp53^FRT/FRT^* Fib^ΔZeb1^;Δp53 and Col6a1-Cre^Tg/+^;*Zeb1^fl/fl^*;Vil-Flp^Tg/+^;*Tp53^FRT/FRT^* Fib^Ctrl^;Δp53 littermates. AOM/p53 tumors were induced by six weekly doses of AOM (10mg/kg, i.p.) in 0.9%NaCl, as described previously ^32, 33^. Tumor growth was monitored by endoscopy as in the AOM/DSS model and animals were sacrificed after 161 days or at any other ethical endpoint.

Orthotopic transplantation of tumor organoids was performed as described ^31^. In brief, AKP or AKP^re^ organoids were mechanically dissociated, seeded in high concentration collagen pads (6.5 mg/ml) and allowed to recover for 48 hours. Collagen pads were transplanted below the *muscularis externa* of the mouse caecum and serosal wounds covered by an anti-adhesion barrier. Recombination of *Zeb1* was induced at day 14 for AKP organoids and at day 4 for AKP^re^ organoids by 400 mg/kg tamoxifen diet (custom order based on Altromin 1824P). Tumor growth was monitored three times a week by palpation and mice were sacrificed when tumors exceeded 1 cm in diameter. For immune checkpoint inhibition, mice were injected i.p. with anti-PD-L1 (Bio X cell, BE0101) or matched control antibodies (Bio X cell, BE0090) at days 14, 21 and 28. To ensure equal treatment of mice, only tumors collected after day 28 were considered for size comparison. For AKP^re^ transplantation, mice treated with control antibodies were analyzed instead of untreated animals to reduce total experimental animal numbers.

### Skin wound healing

Wound closure of a skin excision wound in mice was determined over time for 10 days ^58^. Briefly, circular wounds were applied under sterile conditions on anaesthetized mice into the dorsum after depilation. The dorsal skin was lifted at the midline and punched through two layers of skin using a disposable biopsy punch (6 mm in diameter, Kai medical, Solingen #BP-60F). Wound size was determined at indicated time-points with a digital Caliper and by transferring wound outline to a transparent film for area calculation using ImageJ.

### Organoid culture and genetic engineering

Primary mouse intestinal organoid cultures were established as reported ^59^. Briefly, mouse colons were cut into small fragments, washed with PBS and epithelial crypts were dissociated by incubation with 10 mM EDTA, passed through a 100 µm mash and collected by centrifugation. Colon crypts were seeded at high density in BME (R&D Systems, 3533-010-02) and cultured in Advanced DMEM/F12 (Thermo Fisher Scientific, 12634028) with 20 % Wnt3a conditioned medium, 10 % Noggin conditioned medium, 10 % R-spondin1 conditioned medium, B27 supplement (Thermo Fisher Scientific, 17504-044), 500 mM N-Acetylcysteine (Merck, A9165), 500 µg/mL human EGF (Peprotech, AF-100-15) and 500 µM A83-01 (Tocris, 2939/10). Established organoids were passaged twice a week by mechanical dissociation and reseeding.

For generation of tumor organoids, colons from Lox-Stop-Lox-Kras^G12D^ mice were used ^60^. Both *Apc* and *Tp53* alleles were mutated by co-transfection of Cas9 with the respective sgRNA plasmids. The oncogenic *Kras^LSL.G12D^* allele was recombined by pAC1-Cre (ATCC, 39532) transfection. Modified lines were clonally expanded and successful modification confirmed by Sanger sequencing.

The hCas9 and gRNA_GFP-T2 plasmids were a kind gift from George Church (Mali, et al. 2013, Addgene plasmids #30205 and #41820) and the Apc sgRNA plasmid kindly provided by Hans Clevers ^61^. sgRNA oligonucleotides targeting *Tp53* (sense: 5’-GTTTTAGAGCTAGAAATAGCAAG-3’ antisense: 5’-AGTGAAGCCCTCCGAGTGTCGGTGTTTCGTCCTTTCCACAAGAT-3’) were inserted into gRNA_GFP-T2 plasmid by inverse cloning as published ^61^.

Tumor organoids from established tumors were obtained by enzymatic digestion of primary tumors with 0.1 % Collagenase I (Thermo Fisher Scientific, 17018029), 0.2 % Dispase II (Merck, D4693) and 2 U/mL DNase I (New England Biolabs, M0303L). Cells were passed through a 40 µm mash and seeded at high density in BME.

### Primary colon fibroblast isolation and culture experiments

For fibroblast cultures normal colon fragments after epithelial crypt isolation or minced AOM/DSS tumors were washed in PBS and further incubated with 10 mM EDTA. After 1 hour, tissue fragments were seeded into 10 cm culture plates coated with 0.1 % gelatin (Sigma, G2500) in DMEM (Thermo Fisher Scientific, 31966-047) with 10 % FBS (Sigma, F7524) and Pen/Strep (Thermo Fisher Scientific, 15140-122). After several days, fibroblasts expanded from tissue fragments attached to the plates, which were collected by trypsinization and propagated as regular 2-dimensional cell cultures on gelatin coated plates.

For the collagen contraction assay, 1 x 10^5^ fibroblasts were seeded in 500 µL Collagen I solution (Thermo Fisher Scientific, A1048301) into a 24-well suspension plate. After polymerization of collagen, the collagen disks were detached from the plate and contraction was determined by monitoring the area of the disks over time.

For the barrier function and T cell attraction assays, splenocytes were isolated from OT1 mice ^62^. T cells were stimulated by addition of SIINFEKL peptide (AnaSpec, AS-60193-1) for 4 hours and expanded for 3 days prior to an experiment. For the barrier function assay, 12-well transwells with 8 µm pore size (Greiner, 665638) were coated with 0.1 % gelatin and 2 x 10^5^ fibroblasts were seeded on top of the transwell. After 48 hours, 3 x 10^4^ T cells were added on top of the confluent fibroblast layer and cells in the lower compartment were monitored microscopically. For the T cell attraction assay, 1 x 10^5^ fibroblasts were seeded into a 12-well adhesion culture plate. After 24 hours, a transwell with 3 µm pore size (Corning, CLS3462-48EA) containing 3 x 10^4^ T cells was added to the well and attracted cells in the lower compartment were monitored microscopically.

For inflammatory activation colon fibroblasts were plated in 6-wells and after 24 hours treated with 1 ng/ml murine recombinant IL-1α (Biolegend, 575002) in PBS for the indicated time periods. To ensure optimal timing the medium was replaced by 1 ml of fresh medium 2 hours before treatment and supplemented with 1 ml 2 ng/ml IL-1α containing medium at the starting point. At indicated time points cells were harvested and processed for RNA and protein isolation.

### Histology, immunohistochemistry (IHC) and immunofluorescence labeling (IF)

Tumors of orthotopic transplantation were dissected and fixed in 4 % PFA/PBS overnight at 4°C. For histological and IHC analyses, tumors were sectioned at 3 µm and stained using the Bond-Max device (Leica) (CD4: Cell Signaling Technology, 25229S; CD8a: Synaptic Systems, 361003; FoxP3: R&D Systems, MAB8214; PD-L1: Cell Signaling Technology, 13684S; B220: Thermo Fisher Scientific, 14-0452-82) and the Bond Polymer Refine Detection system (Leica, DS9800). Sections were imaged using an Aperio CS2 digital pathology slide scanner (Leica) and marker expression was quantified with macros in Aperio ImageScope (v12.4).

AOM/DSS and AOM/p53 colons were collected, flushed with PBS and longitudinally opened for imaging and determining tumor sizes and numbers using a Caliper. Colons were mounted as swiss rolls, fixed in 4 % PFA/PBS overnight at 4°C, paraffin embedded, sectioned at 3-4 µm and subjected to haematoxylin/eosin (H&E), IHC or IF staining as described previously ^21^ (cl. Casp. 3: Cell Signaling Technology, 9661S; CD4: Sino Biol. 50134-R001; CD8a: Sino Biol. 50389-T26; FoxP3: ebiosciences, 14-5773-82; PD-L1: LSBio, LS-C19686/144489, B220: Thermo Fisher Scientific, 14-0452-82, anti-rabbit-polymer: DAKO, K4003; anti-rat-HRP: Life Technologies, A18915). After antibody incubations, washing steps and DAB reactions, the slides were counterstained with Mayer’s haematoxylin before dehydration and mounting (Roti®-Histokitt, 6638.2). Analysis and image acquisition was performed using a Leica DM5500B microscope. Scoring of infiltration into AOM/DSS tumors was estimated from the cellularities of infiltrated cell-type marker-positive cells on IHC slides, with none (0), rare (1), few (2), several (3) and many/abundant (4) classification. For anti-ZEB1 immunofluorescence staining of Fib^Ctrl^/Fib^ΔZeb1^ (Col6a1-Cre) mice, cryosections from fresh frozen colon specimens were fixed in 4 % PFA for 10 min, permeabilized for 10 min in 0.25 % Triton X-100/PBS, blocked in 3 % BSA/PBS and incubated with anti-ZEB1 antibodies (Sigma, HPA027524, 1:200 in 3 % BSA/PBS). After washing and incubation with Alexa594-conjugated secondary antibodies (Life Technologies, 1:500 in PBS), DAPI stained sections were mounted (Antifadent AF1, Citifluor). Images were acquired using a Leica DM5500B microscope.

For immunofluorescence staining of tumors from orthotopic transplantation, citrate-based antigen retrieval was applied with deparaffinized sections. Slides were blocked with 20 % goat serum (Merck, G9023) in PBST and stained for 1 hour with primary antibodies (ZEB1: Bethyl, IHC-00419; E-Cadherin: BD Biosciences, 610181; Collagen 6: Abcam, ab182744; αSMA: Thermo Fisher Scientific, 14-9760-82) and for 30 min with secondary Alexa488/647-conjugated antibodies (anti-mouse-/anti-rabbit-IgGs: Thermo Fisher Scientific, A-21202/A-31573). Sections were imaged using an Evos FL microscope and marker expression was quantified using CellProfiler ^63^.

### Picrosirius red staining

For picrosirius red staining, slides were deparaffinized, rehydrated and then incubated in Picrosirius Red solution (abcam, ab246832) for 1 h at RT before washing in 0.5% acidified water, dehydration and mounting (Roti®-Histokitt, 6638.2). Polarized light imaging was done by using Leica DM5500B equipped with a polarization filter to monitor green thinner collagen fibers and yellow-orange bundled fibers by refraction and birefringence. Percentages of total stained areas were analyzed on polarized light images using CellProfiler by RGB conversion, global background-based thresholding for pixel identification, followed by binary image generation of each channel for measuring area occupied by pixels.

### Western blot analysis

Protein extraction and western blotting was carried out as described (Lehmann, et al, 2016, Schuhwerk et al., 2022). Briefly, cells grown in 6 well-plates were lysed in 150 mM NaCl, 50 mM Tris-HCl pH 8.0, 0.5 % Na-Desoxycholate (w/v), 0.1 % SDS (w/v), 1 % NP40 (v/v), 1x complete protease inhibitor (Roche, 4693132001), 1 mM PMSF, 1x PhosSTOP (Roche, 4906837001) for 20 min at 4°C. Protein concentrations were determined by using the BCA Protein Assay (ThermoFisher Scientific, 23225) according to manufacturer’s instructions. Up to 30 µg of protein lysate was separated by SDS-PAGE and transferred to nitrocellulose membranes before antibody incubation (IκBα: Cell Signaling, CS9242; NFκB p65: Cell Signaling, CS8242; p-IκBα Ser32: Cell Signaling, CS2859; p-NFκB p65 Ser536: Cell Signaling, CS3036; ZEB1: Sigma, HPA02752; α-Tubulin: Sigma, T6199; rabbit IgG-HRP: Dianova, 111-035-144; mouse IgG-HRP: Dianova, 115-035-146). Detection was carried out using Western Lightning Plus-ECL (Perkin Elmer, NEL103001EA) and a ChemiDoc^TM^ Imaging System (BioRad).

### Secretome analysis

For secretome analysis, a commercially available secretome array kit was used (R&D systems, ARY028) according to the manufacturer’s instructions. Samples from AOM/DSS CAF supernatants were prepared by surgically detaching adenomas and mincing using scalpels and incubating the tissue in digestion buffer containing 0.05 % Collagenase D (w/v), 0.3 % Dispase II (w/v), 0.05 % DNase I (w/v), 4 % FBS (v/v), in DMEM/F12 medium (ThermoFisher Scientific, 31331028) for 30-45 min at 37°C with constant agitation. 10 ml of cold washing buffer (sterile PBS containing 2 % FBS) was added, and the suspension passed through a 70 µm strainer. After centrifugation and erythrocyte lysis in ACK buffer (150 mM NH_4_Cl, 10 mM KHCO_3_, 0.1 mM EDTA; pH 7.2-7.4) for 2 min at RT, washed and collected cells were resuspended in DMEM/F12/10 % FBS and plated in 12-well plates. Adherent cells were passaged in 1:1 ratios when reaching confluence. After 2-3 passages, the supernatants of individual confluent 6-well vessels were collected and kept on ice. After centrifugation at 1000 g for 5 min at 4°C, supernatants were aliquoted and frozen at −80°C. For the secretome array, samples were added to the equilibrated and blocked membranes for incubation over night at 4°C. After incubation with the detection antibody cocktail for 1 hour, with the Streptavidin-HRP mix for 30 min and the detection mix for 1 min at RT, secretomes were detected by imaging like for western blot. Intensities were normalized to the background of each respective membrane.

### RNA isolation and quantitative reverse transcriptase (qRT-)PCR

Total RNA of cultured cells was isolated using the RNeasy Plus Mini Kit (Qiagen, 74136 and 200-500 ng of total RNA was used to synthesize cDNA using the RevertAid First Strand cDNA Synthesis Kit (Thermo Fisher Scientific, K1622) according to manufacturers’ instructions. Subsequent qRT-PCR of cDNA was performed in triplicates in 384-well plates using primers and Roche universal probe library (UPL) with TaqManTM Universal MasterMix II (Thermo Fisher, 4440044) and LightCycler® 480 II (Roche).

### Single cell RNA sequencing (scRNA-Seq) and analysis

Single cell transcriptomes were generated using a commercially available 384-well plate approach (SORT-Seq2, Single Cell Discoveries, SCD ^64^). To this end, primary tumors were dissected and dissociated into single cells as for establishment of organoid lines from tumors. After erythrocyte lysis in ACK buffer, cells were incubated with Fc Block, anti-CD45-PE/Cy7, anti-CD31-PE, anti-Epcam-eFluor450, a-CD140a-APC and eFluor780 (BD Biosciences, 553142; Biolegend, 103113; Thermo Fisher, 12-0311-82, 48-5791-82, 17-1401-81 and 65-0865-14) in PBS/2 % FCS/2 mM EDTA. Single viable cells (eFluor-) were gated for Epcam^pos^ or CD45^pos^ or double negative cells. Epcam^neg^, CD45^neg^ cells were further gated for CD31 expression and triple negative cells were considered fibroblasts for sequencing. Cells were index-sorted into 384-well capture plates containing 50 µl lysis buffer and barcoded primers covered by 10 µl of mineral oil by flow cytometry using a BD FACSAria Fusion fluorescence activated cell sorter (BD biosciences) and capture plates were sent for paired-end sequencing at SCD (Illumina Nextseq™ 500). Sequences from read 1 were used for assigning reads to cells and libraries, whereas read 2 was aligned to the ensemble transcriptome (genome assembly GRCm38) using *bwa* version 0.7.10 ^65^. The transcript count table was generated by SCD using a custom-written script (https://github.com/anna-alemany/transcriptomics/tree/master/mapandgo). Transcript counts and metadata for all samples were imported and stored as a single-cell experiment object (*SingleCellExperiment* ^66^ version 1.12.0). During quality control, we discarded cells with high mitochondrial content (*isOutlier*, scater ^67^ version 1.18.5) to remove low-quality cells that may have been damaged during processing or may not have been fully captured by the sequencing protocol. Additionally, cells with high expression of *Ptprc* and *Epcam* genes were excluded to avoid contamination by residual immune/ epithelial cells that were insufficiently eliminated by FACS prior to sequencing. After quality control, the samples were normalized by computing the log-transformed normalized expression values across all genes for each cell (*logNormCounts*, scater). Next, all batches were subset to the common features across all samples to enable downstream analysis. To account for the differences in samples due to plates (‘batch effect’), we used the fastMNN algorithm (*correctExperiments*, batchelor ^68^ version 1.6.2). Subsequently, clusters were identified using a graph-based approach (*buildSNNGraph* with k set to 15, scran ^69^ version 1.18.5) and the walktrap algorithm (igraph version 1.2.6). Cluster-specific marker genes were identified using the *findMarkers* function (scran) that uniquely define one cluster against the rest. Trajectory analysis was performed using Slingshot (version 1.8.0) to identify the path traversed by cells through different states. Functional enrichment analysis in Metascape (https://metascape.org/) and Enrichr (https://maayanlab.cloud/Enrichr/) was applied to identify pathways and processes that were enriched in each cluster based on differentially expressed genes (FDR≤0.1; p≤0.05). Furthermore, we compared the expression profile of identified clusters with those previously published for CAF subtypes using the *AddModuleScore* function ^70^ from Seurat ^71^ version 4.0.0. To this end, each cell was assigned a score using the module of genes associated with the ‘published clusters’. A positive score suggested that this module of genes is expressed in a particular cell more than would be expected, given the average expression of this module across the population. A mean score for cluster-specific cells was calculated to obtain scores per cluster for each module of genes associated with ‘published clusters’. Using a reference dataset with known labels, the SingleR ^72^ approach (version 1.4.1) labels new cells from a test dataset based on similarity to the reference. Thereby we assessed similarity between clusters from Fib^ΔZeb1^ and Fib^Ctrl^, where Fib^Ctrl^ was set as a reference. For the scRNA-seq dataset from non-inflammation-driven orthotopic tumors lower quality clusters with indistinct or ambiguous cell-type identities were excluded by further sub-clustering.

### Statistics

Statistical analyses were performed using GraphPad Prism, within the provided R packages or online tools for gene set enrichment analyses. Details are provided in the respective figure legends.

### Data availability

scRNA-seq data will be deposited at GEO to be publicly available as of the date of publication. All original code has been deposited at Zenodo and is publicly available as of the date of publication. Any additional information required to reanalyze the data reported in this paper is available upon request.

## Acknowledgements

We thank Britta Schlund, Eva Bauer, Friederike Gräbner, Tahmineh Darvishi and Petra Dinse for excellent technical assistance. We are grateful to Stefan Stein and Annette Trzmiel (Core Facility Flow Cytometry GSH Frankfurt) for support with all fluorescence-activated cell sorting (FACS) and the animal facilities at FAU Erlangen and GSH Frankfurt for excellent animal care and welfare. We thank all members from the Farin, Stemmler and Brabletz laboratories for critical input and fruitful discussions. This work was supported by the German Research Foundation (FOR2438/P04, TRR305 TP A03, A04, B01 and B07, and BR1399/9-1, BR1399/10-1, BR4145/1-1 and BR4145/2-1), the Wilhelm Sander-Stiftung (2020.039.1; 2019.143.1) and the European Union’s Horizon 2020 research and innovation program under the Marie Skłodowska-Curie grant agreement (No. 861196, PRECODE).

## Extended Data Figures & Legends

**Extended Data Fig. 1.**
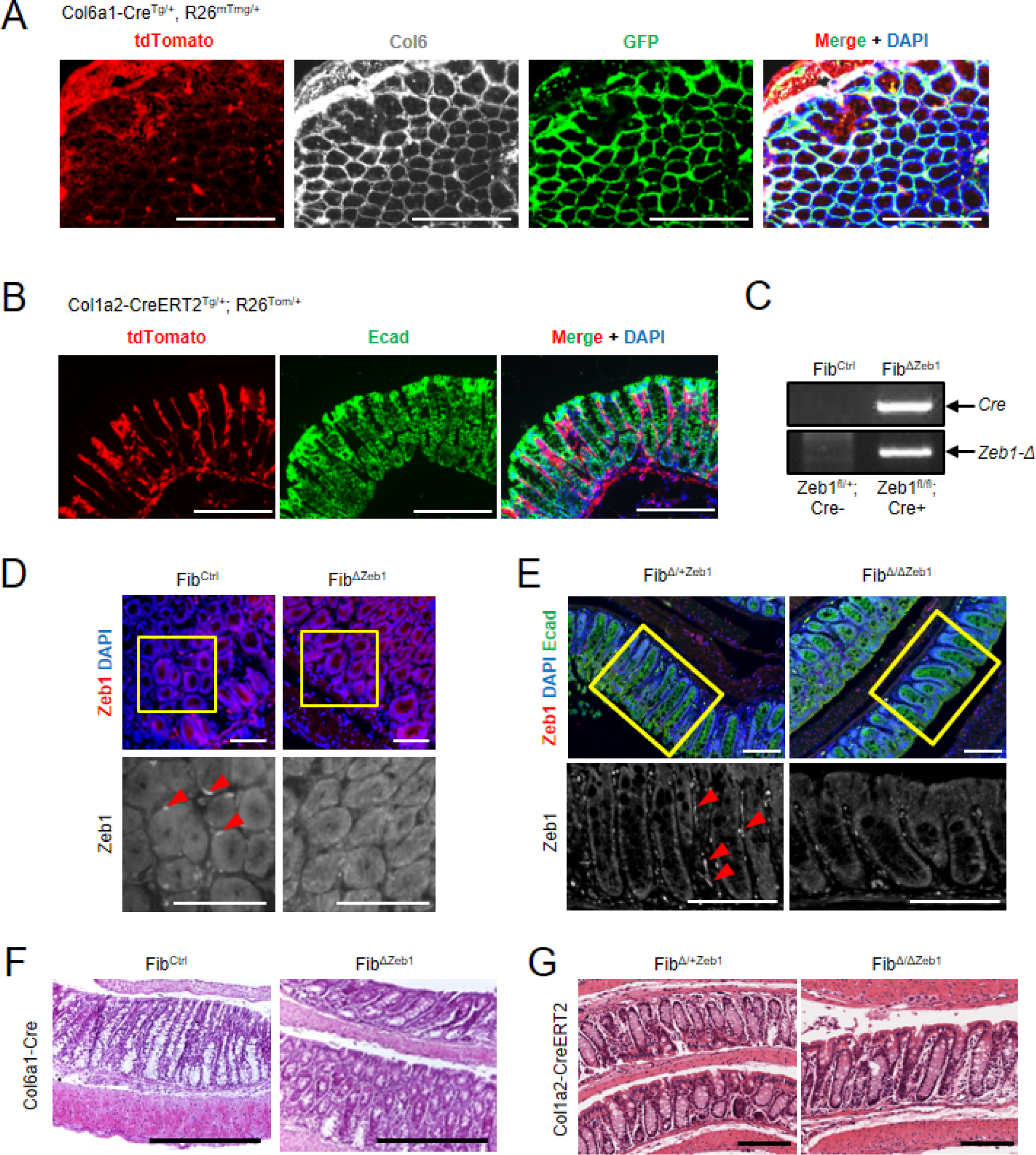
Loss of *Zeb1* in fibroblasts cells does not affect intestinal homeostasis. (A) Col6 immunofluorescence (IF) and DAPI staining on cryosections with tdTomato and GFP fluorescence overlay on colon sections of Col6a1-Cre^Tg/+^;Rosa26-mTmG^Tg/+^ mice. Scale bar: 200 µm. (B) IF of tdTomato, E-cadherin (Ecad) and DAPI on colon sections of Col1a2-CreERT^Tg/+^;Rosa26-tdTomato^Tg/+^ mice. Scale bar: 200 µm. (C) Genotyping of colon DNA confirms presence of Cre transgene and Col6a1-Cre induced recombination of *Zeb1* in indicated mouse genotypes. (D, E) IF of ZEB1 and DAPI staining on colon sections of Fib^Ctrl^ (*Zeb1^fl/fl^*;Col6a1-Cre^+/+^) and Fib^ΔZeb1^ (*Zeb1^fl/fl^*;Col6a1-Cre^Tg/+^) mice (D) and IF of ZEB1, Ecad and DAPI on colon sections of heterozygous Fib^Δ/+Zeb1^ (*Zeb1^fl/+^*;Col1a2-CreERT2^Tg/+^) and Fib^ΔZeb1^ (*Zeb1^fl/fl^*;Col1a2-CreERT2^Tg/+^) tamoxifen-fed mice (E). Bottom rows show magnification of ZEB1 of the marked areas in grayscale. Red arrowheads indicate ZEB1 positive stromal cells which are absent in Fib^ΔZeb1^. Scale bars: 100 µm. (F, G) H&E stainings of colon sections from Fib^Ctrl^ and Fib^ΔZeb1^ mice (F), as well as of Fib^+/ΔZeb1^ and Fib^ΔZeb1^ tamoxifen-treated mice (G). Scale bars: 200 µm.

**Extended Data Fig. 2.**
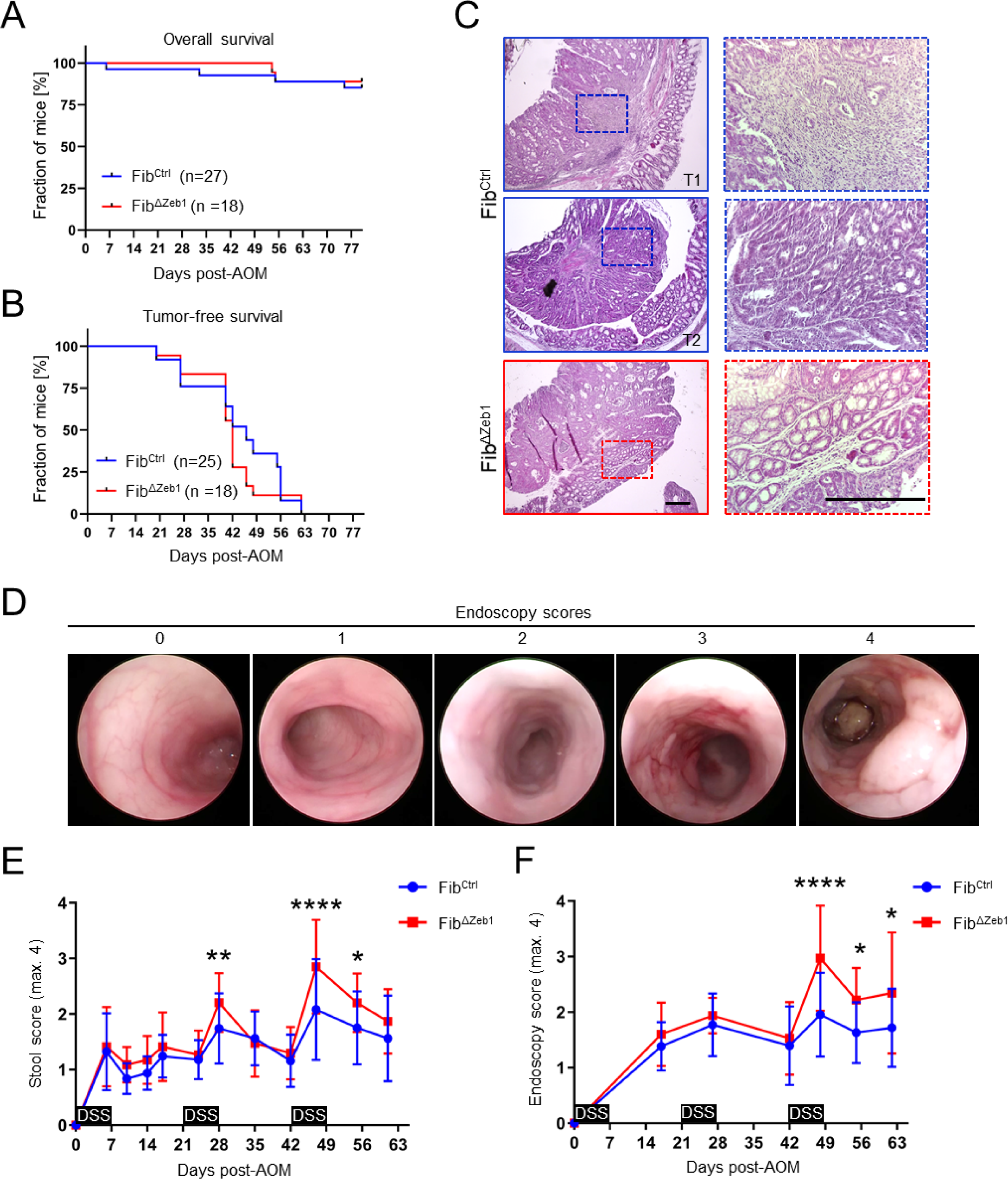
ZEB1 in fibroblasts moderately exacerbates AOM/DSS-induced colitis. (A, B) Kaplan-Meier plots showing overall survival (A) and tumor-free survival (B) of Fib^Ctrl^ and Fib^ΔZeb1^ mice during AOM/DSS tumorigenesis. Numbers of experimental mice are indicated. (C) Representative images of H&E stainings of colon adenoma sections from Fib^Ctrl^ and Fib^ΔZeb1^ mice. A higher magnification of the indicated region is given to the right. Scale bars: 300 µm. (D, E) Representative images during endoscopy correlated with identified endoscopy scores (D) and quantification of intestinal inflammation based on stool scoring (E) and on endoscopy scoring (F, also refer to images in D) in Fib^Ctrl^ and Fib^ΔZeb1^ mice during AOM/DSS tumorigenesis (n=25/17 (E) and n=25/18 (F) for Fib^Ctrl^/Fib^ΔZeb1^, mean ± SD, *: p < 0.05, ****: p < 0.0001, 2way ANOVA).

**Extended Data Fig. 3.**
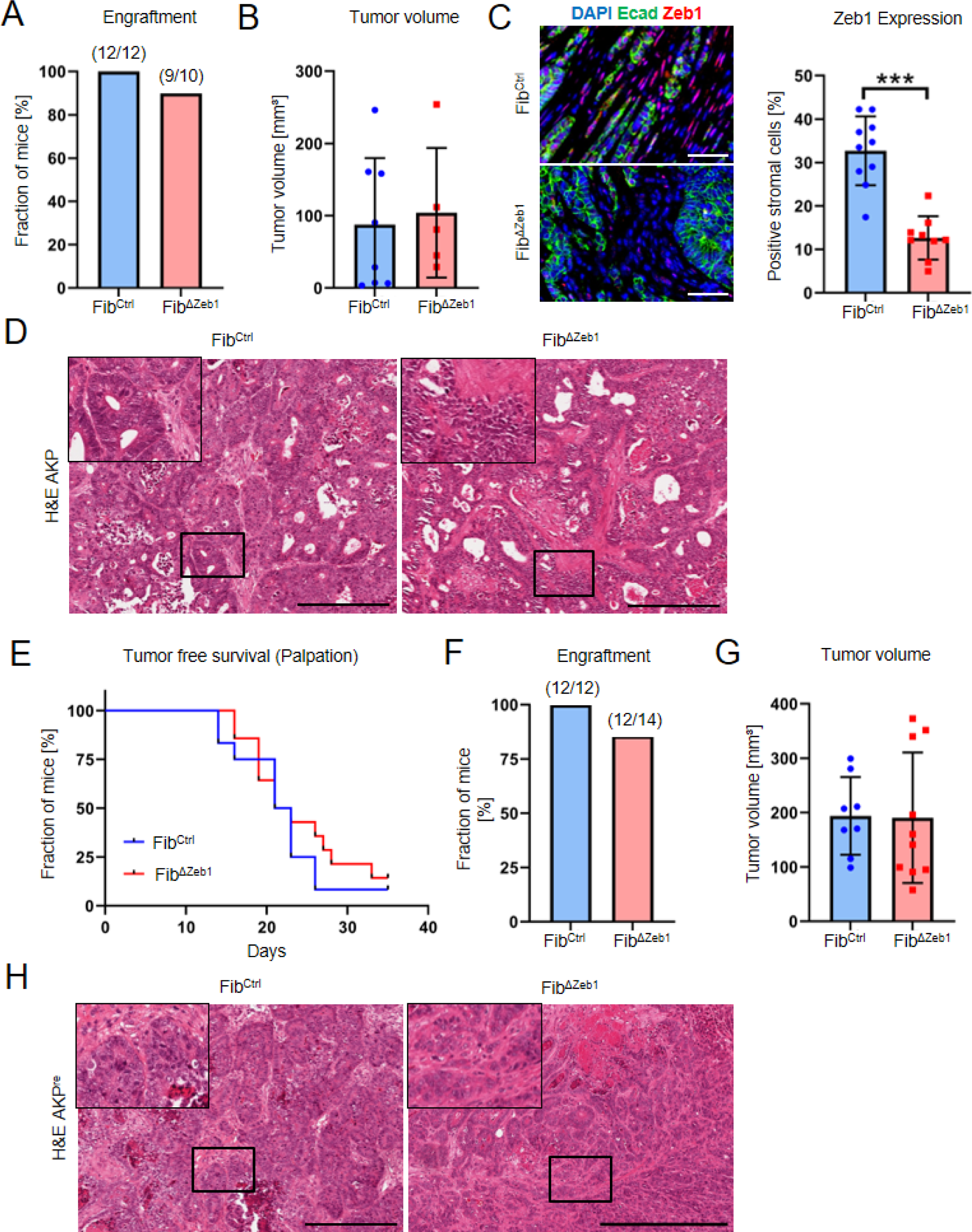
Loss of *Zeb1* in fibroblasts does not affect morphology of primary tumors in the orthotopic transplantation model. (A) Quantification of primary tumor engraftment in Fib^Ctrl^ and Fib^ΔZeb1^ mice after orthotopic transplantation of AKP tumor organoids. Numbers of experimental mice are indicated. (B) Tumor volume after orthotopic transplantation of AKP tumor organoids (n=8/5 for Fib^Ctrl^/Fib^ΔZeb1^, mean ± SD, ns by student’s t-test). (C) Immunofluorescence images and quantification of ZEB1 expression in tumor stroma after orthotopic transplantation of AKP tumor organoids. Scale bar: 50 µm (n=10/9 for Fib^Ctrl^/Fib^ΔZeb1^, mean ± SD, ***: p < 0.001, student’s t-test). (D) Representative H&E stainings of AKP tumor sections. Top left corners show higher magnification of the indicated regions. Scale bars: 1 mm. (E) Analysis of tumor onset after orthotopic transplantation of AKP^re^ tumor organoids (n=12/14 for Fib^Ctrl^/Fib^ΔZeb1^, ns by Mantel-Cox test). (F) Quantification of tumor engraftment after orthotopic transplantation of AKP^re^ tumor organoids. Numbers of experimental mice are indicated. (G) Tumor volume after orthotopic transplantation of AKP^re^ tumor organoids. Only tumors collected after day 28 were included in this analysis (n=8/10 for Fib^Ctrl^/Fib^ΔZeb1^, mean ± SD). (H) H&E staining of representative sections after orthotopic transplantation of AKP^re^ tumor organoids. Top left corners show higher magnification of the indicated regions. Scale bars: 1 mm. All mice with AKP^re^ transplantation were treated with control IgG.

**Extended Data Fig. 4.**
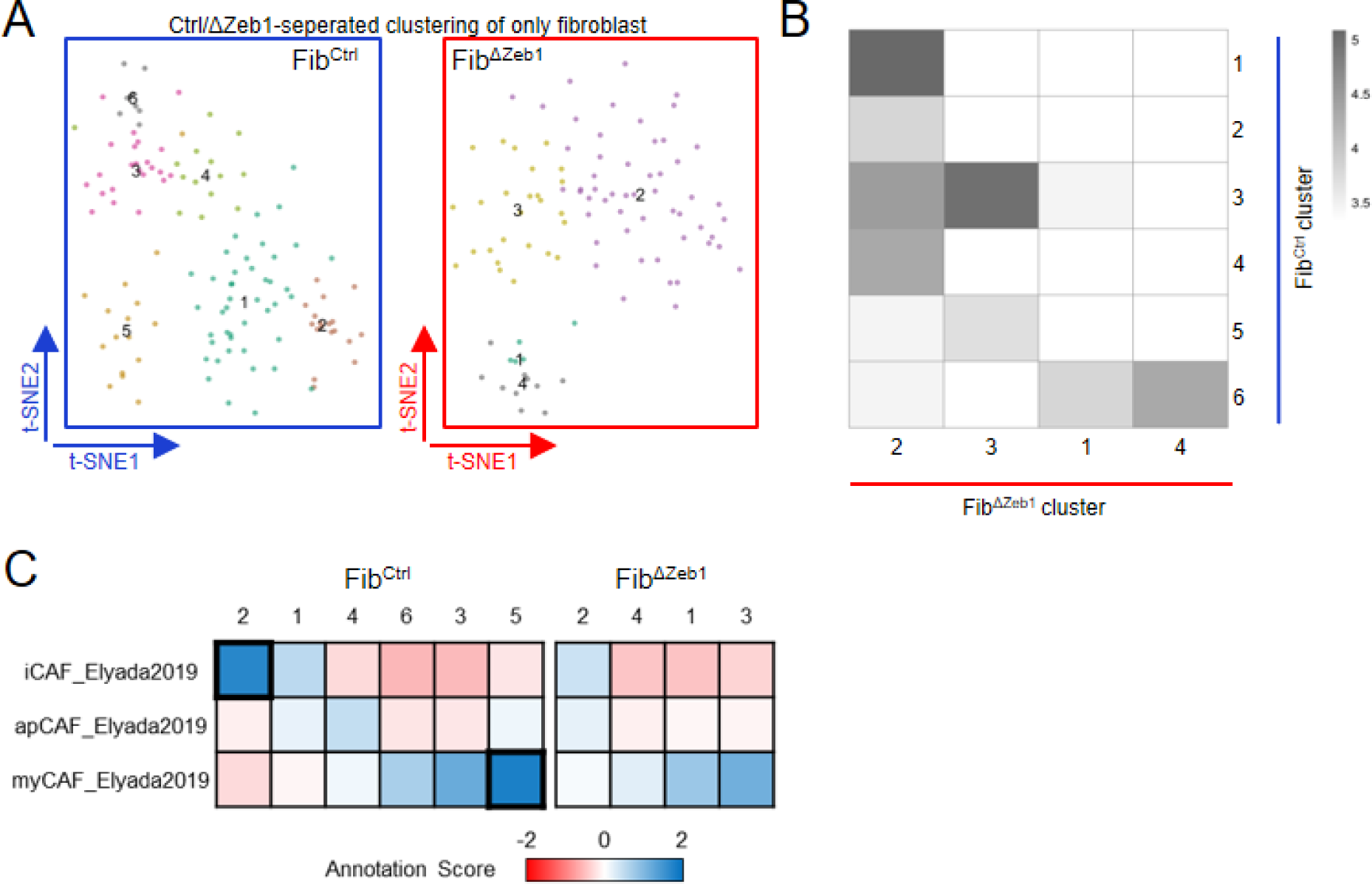
scRNA sequencing analysis of CAFs from the orthotopic model separately shows reduced diversity and impairment of CAF subtype expression profiles in Fib^ΔZeb1^ tumors. (A) Unbiased separated t-SNE sub-clustering of fibroblasts from Fib^Ctrl^ (left) and Fib^ΔZeb1^ mice (right) of the orthotopic model showing less clusters in Fib^ΔZeb1^. (B) Reference-based annotation of fibroblast clusters of Fib^ΔZeb1^ mice using the single-cell gene expression profiles of Fib^Ctrl^ clusters as reference with the heatmap showing the distribution of cells across Fib^ΔZeb1^and Fib^Ctrl^ clusters based on ‘SingleR’ scores ^72^. Color scale in the heatmap shows the log-transformed number of cells across clusters. Note the low similarity of Fib^ΔZeb1^ cells with Fib^Ctrl^ clusters 2 and 5. (C) Heatmap showing the similarity (annotation scores) of gene expression in CAF clusters with published gene sets. Note, the high scores of Fib^Ctrl^ clusters 2 and 5 when compared with ‘iCAF’ and ‘myCAF’ signatures, respectively, and absence in Fib^ΔZeb1^.

**Extended Data Fig. 5.**
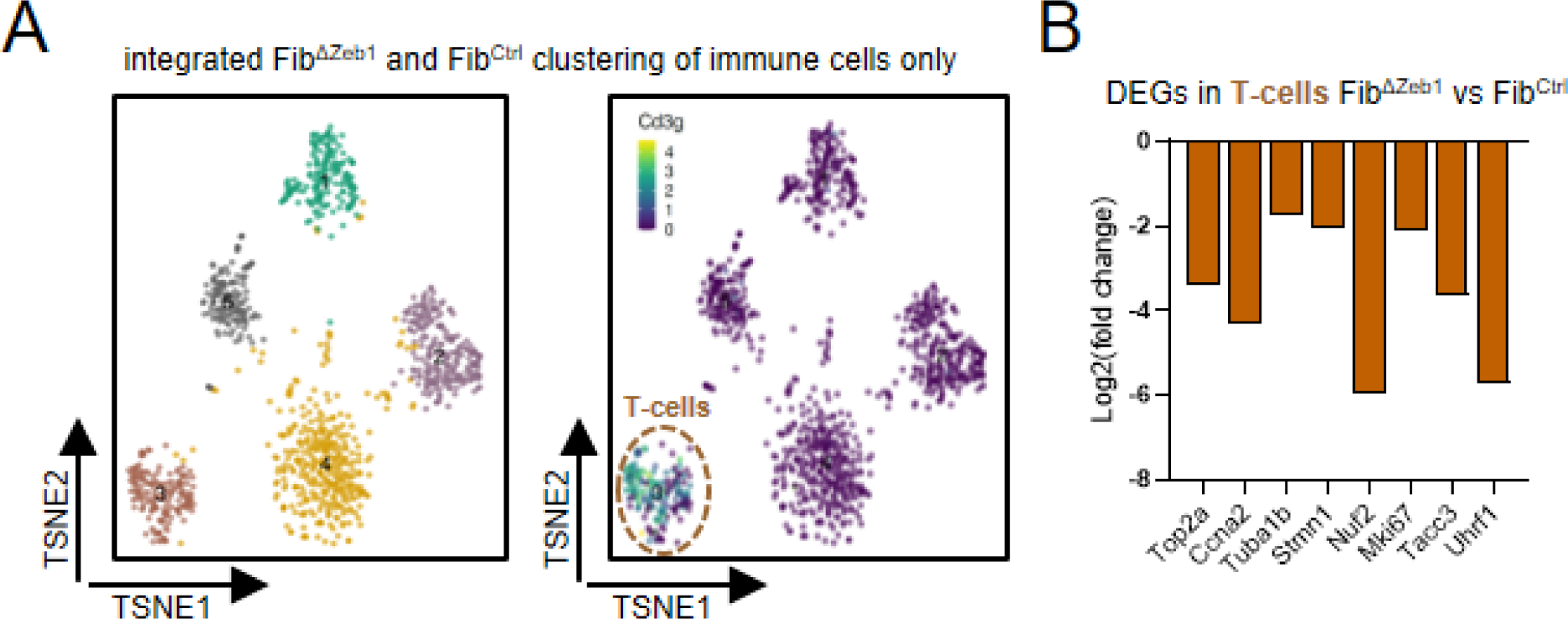
T cells are less proliferative in orthotopic tumors after loss of *Zeb1* in CAFs. (A) t-SNE sub-clustering of immune cells from tumors of Fib^Ctrl^ and Fib^ΔZeb1^ after orthotopic transplantation of AKP organoids (left) and projection of the relative expression level of the representative T cell marker gene *CD3g* as t-SNE feature plot (right), identifying cluster 3 as T cells. (B) Log2-fold change of differentially expressed genes (FDR<0.05) in Fib^ΔZeb1^ versus Fib^Ctrl^ T cells.

**Extended Data Fig. 6.**
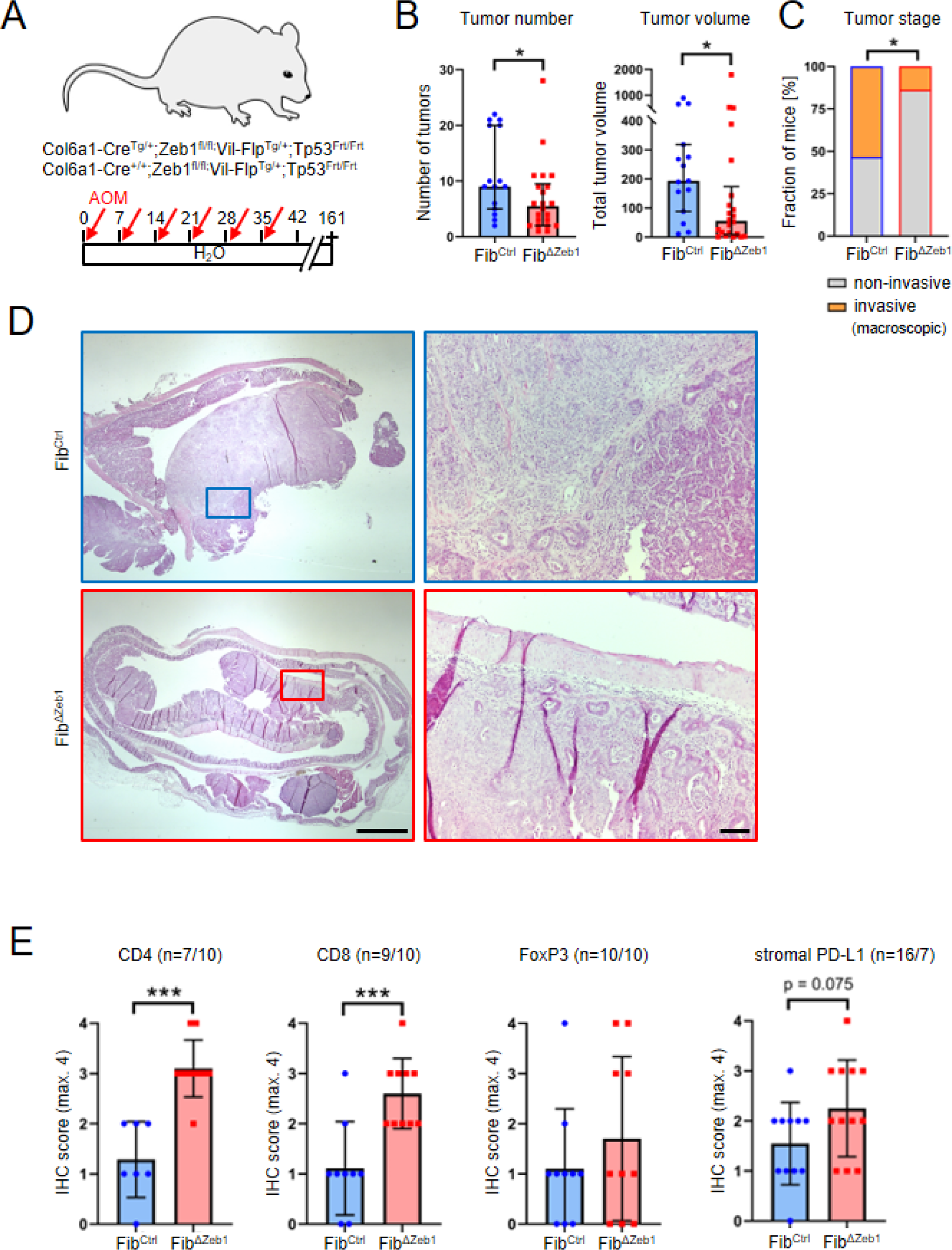
Loss of *Zeb1* in fibroblasts impairs tumor progression and increases T cell infiltration in the invasive non-inflammation driven AOM/p53 model. (A) Schematic representation of the AOM/p53 model. (B) Quantification of numbers and volumes of tumors in the colons of Fib^Ctrl^ and Fib^ΔZeb1^ mice at the endpoint (n=15/22 for Fib^Ctrl^/Fib^ΔZeb1^, mean ± SD, *: p < 0.05, Mann-Whitney U-test). (C) Macroscopic evaluation of the most advanced/progressed tumor per mouse categorizing ‘T4’ as fully invasive (penetrating the muscle) or not (‘T1-T3’) (fraction of mice is given, n=15/22 for Fib^Ctrl^/Fib^ΔZeb1^, *: p < 0.05, Fisher’s exact test). (D) Representative H&E stainings with a higher magnification of the indicated region to the right. Scale bars: 1.5 mm (left) and 100 µm (right). (F) IHC-based quantification of immune cell infiltration and PD-L1 expression of tumors from Fib^Ctrl^ and Fib^ΔZeb1^ mice. Numbers of experimental mice per genotype are indicated (mean ± SD, ***: p < 0.001, student’s t-test).

**Extended Data Fig. 7.**
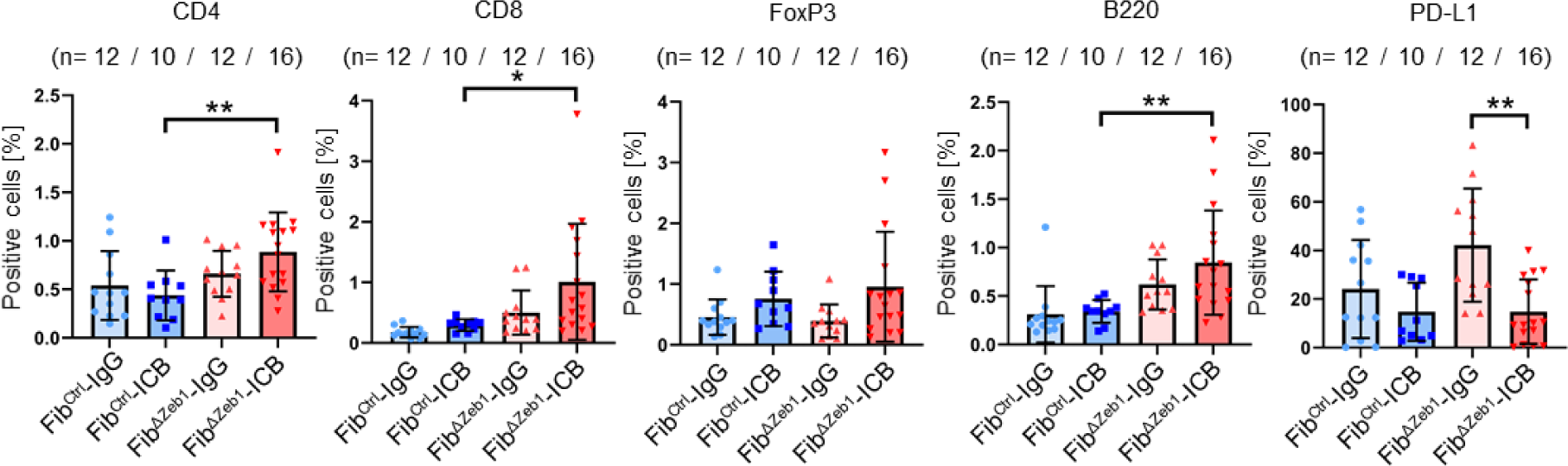
Immune cell infiltration due to loss of *Zeb1* in CAFs is further enhanced by ICB. (A) IHC-based quantification of immune cell infiltration and PD-L1 expression in tumors from Fib^Ctrl^ and Fib^ΔZeb1^ mice after orthotopic transplantation of AKP^re^ organoids and intraperitoneal injection of a-PD-L1 antibodies or control IgGs. Numbers of experimental mice per condition are indicated (mean ± SD, *: p < 0.05, **: p < 0.01, one-way ANOVA) and IgG-treated mice contributed to initial AKP^re^ analysis.

